# An optimized flow cytometry sorting-sequencing workflow reduces storage, sorting and extraction bias in sorted microbial communities

**DOI:** 10.1101/2025.11.13.688239

**Authors:** Valerie Goethals, Nathan Vanwinsen, Fabian Mermans, Sofie De Gand, Nico Boon, Tom Van de Wiele, Kim De Paepe

**Author notes:** Both authors contributed equally to this work (Valerie Goethals, Nathan Vanwinsen). To whom correspondence should be sent.

## Abstract

Fluorescence-Activated Cell Sorting (FACS) followed by metataxonomics is commonly used to probe the composition of microbial subpopulations with a functionality of interest. Metataxonomic analysis of sorted low microbial biomass samples is prone to contamination and bias introduced during sorting and DNA extraction. To improve metataxonomic accuracy, the DNA yield obtained by the DNeasy® PowerSoil® Pro kit was first maximized. FACS-sequencing using the optimized DNA extraction method distorted the composition of sorted microbiota by introducing sheath fluid contaminants, termed the FACSome. Carefully selected controls allowed for FACSome characterization and subsequent *in silico* decontamination of PacBio and Illumina metataxonomic sequencing data. The optimized extraction and decontamination were validated in soil, water, feces, saliva, gut reactor and mock microbial communities. Metataxonomic profiles of non-selectively sorted and unsorted samples derived from these ecosystems did not perfectly match due to relic DNA depletion during FACS. A nuclease-based digest enhanced the depletion of spiked and naturally present relic DNA, which can interfere with function-driven FACS. Since FACS depleted relic DNA, the preservation of intact cells prior to sorting is essential for activity– or viability-targeted FACS. Intact cells were best preserved during long-term –80°C storage of fecal samples that were not amended with glycerol-1X Tris-EDTA cryoprotectant.

## Introduction

Microbial communities perform diverse essential functions in the biosphere. These functions are encoded by the genetic blueprint of individual microorganisms in the community. Functional annotation of this genetic information is non-trivial, yet crucial to understand, monitor, and steer microbial ecosystem functions.

Since individual microorganisms are difficult to analyze, functional genomics traditionally rely on high-throughput bulk meta-omics methods to annotate gene functions. Shotgun metagenomics is the standard method to assess the functional potential of a microbial community based on high-throughput DNA sequencing. The sequenced DNA can contain relic extracellular DNA released from dead microorganisms and DNA from metabolically inactive dormant cells that do not contribute to ecosystem functionality (Amato et al., 2014; Nielsen et al., 2007). In contrast to DNA, mRNA is short-lived and metatranscriptomics high-throughput mRNA sequencing offers a less biased assessment of functionality (Moran et al., 2013). While mRNA transcripts drive ecosystem functions, proteins synthesized by transcript translation perform the functions. Protein and mRNA levels are uncorrelated in single cells and in dynamic microbial communities due to the mismatched half-lives of mRNA (∼ 5 minutes) and proteins (∼ 20 minutes) (Moran et al., 2013). Consequently, metatranscriptomics effectively captures functional shifts, whereas metaproteomics provides a functional snapshot. The end products of chemical reactions catalyzed by proteins still depend on substrate availability and post-translational regulation of protein activity (Moran et al., 2013). End product measurement through metabolomics, thus, delivers the most accurate and direct read-out of microbial community functionality. The identified metabolites are, however, not uniquely linked to a specific microorganism or functionality and can derive from various species, genetic pathways and substrates. Therefore, metabolomics analyses are often integrated with metagenomics. Integration with metagenomics enables pairing of functional and taxonomic information through computationally intensive *de novo* assembly, binning and annotation of functional and taxonomic marker genes (Pavlopoulos et al., 2023). Annotation relies on homology searching against reference databases which are incomplete resulting in microbial functional dark matter (Pavlopoulos et al., 2023). This limitation extends to all meta-omics techniques. Meta-omics workflows are, furthermore, disruptive and microorganisms with functionalities of interest cannot be acquired in a lab environment (Lim et al., 2024). The isolation of targeted microorganisms with functionalities of interest from complex communities through culturomics is low-throughput, laborious and hampered by a large fraction (> 99 %) of viable but nonculturable microorganisms (Lloyd et al., 2018). Prolonged cultivation and adaptive evolution in laboratory conditions can, moreover, lead to loss/alteration-of-function mutations (Martínez & Lang, 2023).

Fluorescence-activated cell sorting (FACS) is a non-disruptive, culture-independent, high-throughput, single-cell technique that can be used to gain functional insights in microbial ecosystems in combination with labelling, viability staining or autofluorescence (Lambrecht et al., 2017; Minnebo et al., 2024; Rinke et al., 2014). Function-targeted labelling can be achieved with fluorescent substrates, DNA, rRNA or mRNA probes (flow cytometry fluorescence *in situ* hybridization, FLOW-FISH) and proteins (antibodies/*in vivo* reporter labeling/bioorthogonal non-canonical amino acid tagging) (Batani et al., 2019; Baxter et al., 2017; Ciolli Mattioli & Avraham, 2024; Mattelin et al., 2024; Mermans et al., 2023, 2025). Sorted fluorescent targeted microbial cells or subpopulations with a functionality of interest can be further analyzed through sequencing or culturing when non-destructive labeling is used. Sequencing is the method of choice to screen diverse subsamples with minimal disturbance and effort. Single-cell sequencing and genomics require a HEPA-filtered cleanroom and robotic liquid handlers to avoid contamination. Such specialized equipment is not available in standard microbiology research facilities, whereas 16S rRNA gene amplicon sequencing of low biomass microbial subpopulations is routinely used in microbial ecology (Mattelin et al., 2024; Mermans et al., 2025; Stepanauskas et al., 2017). FACS is, moreover, widely accessible and applied, in contrast to Raman-activated cell sorting, which is an alternative low-throughput method that is used together with stable isotope probing or FISH to single out microorganisms that perform functions of interest (Alcolombri et al., 2022; Ge et al., 2022). Despite the broad use and applicability (Deehan et al., 2022; Riva et al., 2020, 2023), function-driven FACS followed by 16S rRNA gene amplicon sequencing and metataxonomic analysis is prone to contamination and biases which have not been evaluated.

We therefore identified and removed contamination and bias introduced during sample storage, FACS, low biomass DNA extraction and sequencing based on a mock community standard, high biomass references and sheath fluid and buffer blanks.

Sample storage is common for environmental and clinical samples. Sample storage must preserve cell shapes as a minimal requirement since FACS requires that fluorescent labels are concentrated on or within targeted cells and unbound labels are often washed away to improve the signal to noise ratio (Batani et al., 2019; Deehan et al., 2022). Function-targeted labelling may additionally require a cultivation step and thus viability (Batani et al., 2019; Deehan et al., 2022; Riva et al., 2020). Since there is no consensus storage protocol to preserve microbial viability and integrity (Achá et al., 2005; Aguirre et al., 2015; Deschamps et al., 2020; Kerckhof, & Boon, 2014), we compared murine and human fecal sample storage at –80 °C without cryoprotectants and in 5 % glycerol – 1X Tris-EDTA buffer which was previously used to store marine and fecal samples prior to function-driven FACS followed by single-cell genomics (Lyalina et al., 2021; Stepanauskas et al., 2017).

FACS subsamples the microbial community resulting in a low biomass targeted subpopulation which is prone to contamination (Weyrich et al., 2019). Contamination can occur during sample handling, labelling, FACS, DNA extraction and sequencing. Contamination can originate from human microbiota (e.g., the skin bacterium *Cutibacterium*), the lab environment and commercial reagents. Contamination of reagents in DNA extraction and sequencing kits is well described and known as the kitome (Stinson et al., 2019). In contrast, sheath fluid used for hydrodynamic focusing in FACS represents an unexplored source of contamination. Besides, relic DNA, which constitutes up to 30 – 80 % of the total bacterial DNA pool in soil, sediment, water and mammalian gut ecosystems, is detected through sequencing and can thus interfere with function-driven FACS if it co-exists in sorted droplets with targeted cells (Lennon et al., 2018; Veal et al., 2000). We evaluated relic DNA contamination with a spike-in experiment and tested magnetic beads or enzymatic treatment to remove relic DNA prior to FACS-sequencing (Czurda et al., 2016; Stinson et al., 2019). We also identified human-derived, laboratory and FACS contamination in short-read Illumina and full-length PacBio 16S rRNA gene sequencing data.

The low microbial biomass in sorted samples, consisting of 10^5^ cells mL^-1^ theoretically corresponding to 0.5 ng DNA, presents challenges for DNA extraction and sequencing. Most library preparation kits for 16S rRNA gene sequencing require higher DNA inputs (Illumina Nextera XT: 1 ng; PacBio: 1 ng; Oxford Nanopore: 10 ng). We selected the best-performing DNA extraction protocol described by Demkina et al. (2023) and optimized bead beating frequency and sample input volume to maximize DNA yields from the sorted low microbial biomass samples.

The optimized DNA extraction protocol was used to characterize and minimize biases in a FACS-sequencing workflow that was validated with sorted microbial subpopulations from feces, saliva, the Simulator of the Human Intestinal Microbial Ecosystem (SHIME) gut reactor, soil and water ecosystems.

## Experimental procedures

### 1 Experimental set-up

#### 1.1 Optimizing DNA extraction for low microbial biomass samples with the DNeasy® PowerSoil® Pro kit

The DNeasy® PowerSoil® Pro kit (Qiagen, Venlo, The Netherlands) showed the highest DNA extraction yield in an initial screening in our lab compared to Chelex® (Bio-Rad Laboratories Inc., Hercules, CA, USA), phenol-isopropanol (Geirnaert et al., 2015) and phenol-ethanol DNA extraction (data not shown) and in a study by Demkina et al. (2023) that compared multiple commercially available DNA extraction kits. To further improve the DNA extraction yield in sorted low microbial biomass samples, we evaluated three adaptations to the PowerSoil® Pro kit that varied in sample input volume and bead beating frequency. The standard protocol uses 250 µg sample and performs two cycles of 30 seconds bead beating at 2,000 rpm followed by 30 seconds cooldown to mechanically disrupt cells. A sample input volume of 250 µl and the standard bead beating frequency were compared with an increased sample input volume of 500 µL with standard bead beating, a sample volume of 250 µl with 5 cycles of intensified bead beating at 4,000 rpm for 15 seconds followed by 45 seconds cooldown and an increased 500 µL sample input volume combined with intensified 4,000 rpm bead beating. These adaptations aside, the standard DNeasy® PowerSoil® Pro Kit protocol was followed. Bead beating was carried out with the PowerLyzer® (Qiagen, Venlo, The Netherlands) instrument.

The PowerSoil® protocol variations were evaluated by extracting Gram-negative *Escherichia coli* and Gram-positive *Bacillus subtilis* pure cultures in technical triplicate (n=3). A Gram-negative and Gram-positive pure culture were used to account for the fact that microbial cell wall composition affects cell lysis and DNA extraction efficiency (Atwood, 2016; Fujimoto et al., 2004). *Escherichia coli* str. K-12 substr. MG1655 (NCBI:txid511145; 16S rRNA gene copy number: 7) and *Bacillus subtilis* subsp. *subtilis* strain 168 (NCBI:txid224308; 16S rRNA gene copy number: 10) were grown in LB medium (Carl Roth GmbH, Karlsruhe, Germany) for 19 hours at 37 °C until the cultures reached exponential phase. Cell concentrations of the exponential phase cultures were measured with flow cytometry (2.3) and adjusted to 10^5^ cells mL^-1^ to mimic sorted low biomass samples and to 10^8^ cells mL^-1^ to serve as a high biomass reference. The DNA yield obtained with the different PowerSoil® extraction protocols was analyzed with qPCR (2.1) and QuantiFluor® (2.2) in technical triplicate (n=3).

The optimized extraction protocol using 500 µL sample input volume and five cycles of 15 seconds bead beating at 4,000 rpm with 45 seconds cooldown was applied in other experiments (1.2, 1.3, 1.4 and 1.5). Extracted DNA was stored at –20 °C prior to analysis.

#### 1.2 Validating DNA extraction with the optimized DNeasy® PowerSoil® Pro protocol for low microbial biomass samples

The optimized DNeasy® PowerSoil® Pro extraction protocol (1.1) was validated by comparing the microbial community composition in DNA extracts of diluted low biomass and high biomass reference samples from a broad range of phylogenetically diverse microbial ecosystems. This approach captures a larger diversity in cell wall characteristics that may interfere with DNA extraction compared to the previously used *Escherichia coli* and *Bacillus subtilis* pure cultures (1.1).

The evaluated microbial ecosystems consisted of a mock community standard, human feces, saliva, the Simulator of the Human Intestinal Microbial Ecosystem (SHIME) gut reactor, soil and water. Samples were collected, purified to isolate the microbiota and remove particulate matter and stored or immediately analyzed as documented below. Stored or fresh microbiota samples were filtered with a 20 µm syringe filter (BD Biosciences, Erembodegem, Belgium) and stained with SYBR® Green I for flow cytometry total cell counting (2.3) and diluted with anaerobic phosphate buffered saline (aPBS) to 10^5^ cells mL^-1^ in low biomass samples and to 10^8^ cells mL^-1^ in high biomass references. Diluted and high biomass reference samples were extracted with the optimized DNeasy® PowerSoil® Pro kit DNA extraction protocol (1.1). DNA extracts were sequenced with PacBio full-length 16S rRNA gene sequencing (2.4) and Illumina V3-V4 amplicon 16S rRNA gene sequencing (2.5). The validity of the optimized DNeasy® PowerSoil® Pro protocol was assessed by determining microbial community similarity (3.5) in low biomass DNA extractions based on the theoretical composition of the mock or high biomass reference samples.

We used the commercially available ZymoBIOMICS Microbial Community Standard D6300 (Zymo Research Corporation, Irvine, CA, USA) to assess DNA extraction biases. The mock community standard contains inactivated cells of three easy-to-lyse Gram-negative bacteria (*Escherichia coli*, *Salmonella enterica*, *Pseudomonas aeruginosa*) and five tough-to-lyse Gram-positive bacteria (*Listeria monocytogenes*, *Lactobacillus fermentum*, *Enterococcus faecalis*, *Staphylococcus aureus*, *Bacillus subtilis*) in DNA/RNA Shield™. The mock community standard was stored at –80 °C.

Human feces was self-collected in an airtight bucket containing an Anaerogen sachet (Thermo Fisher Scientific Inc., Merelbeke-Melle, Belgium) by an anonymous healthy donor, who provided written informed consent. Research with human fecal material was approved by the Ethical Committee of the Ghent University hospital (B670201836318). A 20 % (w/v) fecal slurry was prepared by homogenizing feces in aPBS with a stomacher LabBlender 400 (Led techno, Heusden-Zolder, Belgium). The aPBS supernatant with suspended fecal bacteria was collected after centrifugation at 500 x *g* for 2 minutes (Eppendorf Centrifuge 5430, Novolab, Geraardsbergen, Belgium). The aPBS solution consisted of 8.8 g·L^-1^ _K2HPO4_ (Carl Roth GmbH, Karlsruhe, Germany), 6.8 g L^-1^ KH_2_PO_4_ (Carl Roth GmbH, Karlsruhe, Germany), and 1 g L^-1^ C_2_H_3_O_2_SNa (Merck, Darmstadt, Germany). The aPBS buffer was sparged for 15 minutes and subsequently flushed for 15 cycles with N_2_ gas (Air Liquide, Paris, France) to remove dissolved and headspace oxygen prior to autoclaving for 40 minutes at 121 °C (HMC Europe, Tüßling, Germany).

Saliva was self-collected by an anonymous healthy donor by active drooling in a sterile Falcon tube after overnight fasting. The saliva sample was homogenized by vortexing twice for 15 seconds. An extra 5 seconds vortexing step was included prior to downstream pipetting to avoid inaccuracies due to the high sample viscosity. The saliva sample was immediately analyzed.

A gut reactor sample was obtained from a proximal colon vessel from the Simulator of the Human Intestinal Microbial Ecosystem (SHIME; Van de Wiele et al., 2015) from a previous experiment (11/2022). Sampling was performed as described by Minnebo et al (2023). The sample was stored at –80 °C. Prior to flow cytometry counting and FACS, the sample was thawed at 38°C and diluted 1:10 (v/v) in aPBS.

Garden soil (1 g) was collected at Campus Coupure (Ghent, Belgium) and suspended in 10 mL aPBS. The suspension was vortexed twice for 10 seconds, sonicated (Elmasonic S 30, Elma Ultrasonic, Singen, Germany) for 10 minutes and centrifuged at 500 x *g* for 3 minutes to extract the soil microbial biomass in the supernatant. The soil sample was immediately analyzed.

Water was sampled from the Coupure river (51°03’10’N 3°42’34’E, Ghent, Belgium) and immediately analyzed without purification.

#### 1.3 Identifying and removing contaminants introduced during FACS-sequencing

Contaminants introduced by FACS-sequencing were identified in sorted low biomass samples of a mock community in triplicate (n=3) and in sorted human feces, saliva, SHIME gut reactor, soil and water microbial ecosystems in duplicate (n=2). High biomass (10^8^ cells mL^-1^) references, samples diluted to 10^5^ cells mL^-1^ before DNA extraction (1.2), DNA extracts diluted to a concentration corresponding to 10^5^ cells mL^-1^, as well as, aPBS and FACS sheath fluid blanks were included to track the source of contamination.

Sampling and sample processing of the mock community and diverse microbial ecosystems were conducted as described in 1.2. The aPBS was prepared as described in 1.2. Commercially available BD FACSFlow^TM^ sheath fluid (BD Biosciences, Erembodegem, Belgium) was used for hydrodynamic focusing (2.3) and sorting and consisted of sodium chloride, disodium hydrogen phosphate, 2-phenoxyethanol, sodium fluoride, potassium chloride and potassium dihydrogen orthophosphate. FACS sheath fluid was sampled in the fluidics chamber in duplicate (n=2). All samples were diluted in aPBS to an appropriate range for FACS (10^5^ – 10^6^ cells mL^-1^ corresponding to 6,000 events s^-1^) based on the obtained flow cytometry cell counts (2.3). Samples were stained with SYBR® Green I (2.3) and sorted (2.3). Samples were frozen at –20 °C prior to DNA extraction.

Sorted, diluted and high biomass reference samples and blanks were extracted with the optimized DNeasy® PowerSoil® Pro kit DNA extraction protocol (1.1). DNA extracts were sequenced with Illumina V3-V4 amplicon 16S rRNA gene sequencing and PacBio full-length 16S rRNA gene sequencing (2.4). Contamination was identified by comparing the absence/presence of taxa in the sorted samples *versus* blanks, the theoretical mock composition or high biomass reference samples. The FACS-sequencing data was decontaminated (3.4), after which microbial community similarity in sorted low biomass samples was evaluated based on mock or high biomass reference samples (3.5).

#### 1.4 Assessing and eliminating extracellular relic DNA contamination during <u>sorting with a DNA spike-in</u>

FACS is based on the separation of droplets with targeted microbial cells from the non-targeted microbial cell subpopulation. While sorting in the ‘Purity’ precision mode ensured that only droplets containing microbial cells of interest were sorted, relic DNA can contaminate the sorted stream. As such, extracellular relic DNA can bias activity– or function-targeted FACS-sequencing. In order to evaluate contamination of the targeted microbial subpopulation with relic DNA, the partitioning of an extracellular DNA spike-in added to a mock community sample into a cell and noise fraction was quantified through PacBio full-length 16S rRNA gene sequencing.

Extracellular DNA was obtained through DNA extraction (1.1) from a stationary culture of *Veillonella atypica* (Rogosa 1965) (basonym: *Veillonella parvula* subsp. *atypica* Rogosa 1965;NCBI txid39777; 16S rRNA gene copy number: 4) in anaerobic BHI3 medium (Carl Roth GmbH, Karlsruhe, Germany). Strain identity was confirmed with Sanger sequencing (2.6).

*V. atypica* DNA extract (3.5 ng) was spiked into 10^7^ cells mL^-1^ ZymoBIOMICS Microbial Community Standard D6300 (41.03 ng; Zymo Research Corporation, Irvine, CA, USA). The ratio of spiked *V. atypica* (6,059,807 16S rRNA gene copies) to mock (58,800,000 16S rRNA gene copies) 16S rRNA gene copies amounted ∼10 %.

An enzymatic Benzonase® Nuclease treatment was tested to remove spiked extracellular *V. atypica* DNA prior to FACS. Thereto, 10^8^ cells mL^-1^ ZymoBIOMICS Microbial Community Standard D6300 were spiked with 35 ng mL^-1^ *V. atypica* DNA extract and treated with 2 mM MgCl_2_ (Carl Roth GmbH, Karlsruhe, Germany) and 4 kU mL^-1^ Benzonase® Nuclease (Merck, Darmstadt, Germany) at 37 °C for two hours. The reaction was stopped by diluting the sample 1:10 (v/v) in 1X PBS prior to FACS (2.3).

Alternatively, 1.8X AMPure XP magnetic beads (Beckman Coulter Life Sciences, Brea, CA, USA) were thoroughly mixed through pipetting with 350 ng mL^-1^ spiked *V. atypica* DNA in 10^9^ ZymoBIOMICS Microbial Community Standard D6300 cells mL^-1^. Following a three minute incubation, beads were removed in a magnet rack for two minutes. Samples were diluted 1:100 (v/v) before FACS (2.3).

A blank sample consisting of aPBS with DNA spike-in (3.5 ng mL^-1^) was included. All samples were sorted with FACS after cell staining (2.3) and extracted with the optimized DNeasy® PowerSoil® Pro kit DNA extraction protocol (1.1). DNA extracts were sequenced with PacBio full-length 16S rRNA gene sequencing (2.4). Contamination was identified by comparing the absence/presence of taxa in the sorted samples *versus* blanks and the theoretical mock composition. The FACS-sequencing data was decontaminated (3.4), after which *V. atypica* relic DNA contamination and community similarity were evaluated in sorted low biomass samples based on the theoretical mock composition (3.5).

#### 1.5 Optimizing storage conditions of human and murine fecal samples

Microbial cell integrity is imperative for function-driven FACS. We investigated the effect of sample storage on cell integrity and FACS-sequencing (2.3) in human and murine fecal samples stored at –80°C for one month with and without 5 % (v/v) glycerol – 1X Tris-EDTA (glycerol-TE) cryoprotectant. Glycerol-TE was prepared by diluting a 20X TE stock solution in aPBS. Human feces were collected from a healthy individual with informed consent (1.2). Murine feces from five 10-week-old healthy male C57BL/6J mice housed in the same cage were collected by the Department of Internal Medicine and Pediatrics (Ghent University, Ghent, Belgium).

Human fecal pellets weighing approximately 1 g and murine fecal pellets weighing 28.07 ± 8.33 mg were freshly analyzed, stored at –80 °C or suspended in glycerol-TE (Carl Roth GmbH, Karlsruhe, Germany; VWR, Radnor, PA, USA) and stored at –80°C, all in duplicate (n=2). Human and murine fecal pellets were suspended with a pellet pestle (Merck, Darmstadt, Germany) followed by vortex mixing in 5 mL, respectively 0.5 mL glycerol-TE. After storage, the slurry was centrifuged at 5,000 x *g* for 10 minutes and the resulting pellet was redissolved in aPBS. The thawed directly frozen feces were suspended with a pellet pestle in aPBS.

All samples were sorted with FACS after viability staining (2.3). Damaged and dead cells with compromised membrane integrity were stained with Propidium Iodide and all cells were counterstained with SYBR® Green I. All samples were extracted with the optimized DNeasy® PowerSoil® Pro DNA extraction protocol (1.1). DNA extracts were sequenced with PacBio full-length 16S rRNA gene sequencing (2.4). Contamination was identified by comparing the absence/presence of taxa in the sorted samples *versus* high biomass references. The FACS-sequencing data was decontaminated (3.4), after which community similarity and significant differences between the sorted fresh and stored samples were determined (3.5).

### 2 Sample analysis

#### 2.1 Real-time quantitative polymerase chain reaction

The 16S rRNA gene copy number was determined in technical triplicate (n=3) in DNA extracts of pure cultures (1.1) with real-time quantitative Polymerase Chain Reaction (qPCR) to evaluate DNA extraction efficacy. The qPCR reaction volume of 20 μL consisted of 10 μL 2x iTaq universal SYBR Green supermix (Bio-Rad Laboratories, Hercules, CA, USA), 2 µL DNA template, 0.8 µL of a 10 µM stock of the forward (338F 5’-ACTCCTACGGGAGGCAGCAG) and reverse primer (518R 5’-ATTACCGCGGCTGCTGG) (Eurogentec, Serraing, Belgium), and 6.4 µL molecular biology-grade water (Labconsult, Ghent, Belgium). Thermal cycling was performed with a QuantStudio^TM^ 3 real-time PCR system (Thermo Fisher Scientific, Merelbeke-Melle, Belgium) in 96 well plates (Thermo Scientific, Fisher Scientific Belgium, Merelbeke-Melle, Belgium). The qPCR amplification protocol consisted of an initial 2 minute denaturation step at 95 °C followed by 40 cycles of 15 seconds denaturation at 95 °C and 1 minute combined annealing/extension at 60 °C.

The 16S rRNA gene copy number was quantified using a standard curve of tenfold diluted gBlocks 16S rRNA gene fragments of *Bifidobacterium breve* JCM 7019 (250 bp: 338F – 518R, Integrated DNA Technologies, Coralville, IA, US) ranging between 10 and 10^8^ copies µL^-1^ (De Pessemier et al., 2025). A standard was included in triplicate in every qPCR assay to ensure that the qPCR efficiency ranged between 85 and 115 %. A melting curve analysis confirmed qPCR primer specificity (60-95 °C, ΔT per 15 minutes = 0.3 °C). All standards and samples displayed a single peak at a melting temperature ranging between 85-90 °C corresponding to the amplified 16S rRNA gene fragment. Data was further analyzed in R version 4.5.0 (3.1).

#### 2.2 Fluorometric DNA quantification

DNA was quantified in technical triplicate (n=3) in DNA extracts of pure cultures (1.1) by a fluorescence assay with the QuantiFluor® dsDNA System (Promega, Madison, WI, US) and GloMax®-Multi+ Detection System (Promega, Madison, WI, US).

#### 2.3 Flow cytometry cell counting and FACS

Total microbial cell counts were measured in pure cultures (1.1), in samples from a mock community, human feces, saliva, the Simulator of the Human Intestinal Microbial Ecosystem (SHIME) gut reactor, soil and water microbial ecosystems (1.2, 1.3, 1.4) and in fresh and stored human and murine fecal suspensions (1.5) according to Van Nevel et al (2013). Samples were filtered with a 20 µm filter (BD Biosciences, Erembodegem, Belgium), diluted with 0.2 μm (Macherey-Nagel, Anderlecht, Belgium) filter-sterilized aPBS, and incubated with a staining mix (1 % v/v) for 20 minutes at 37 °C. The total cell staining mix (1.1, 1.2, 1.3, 1.4) contained SYBR® Green I nucleic acid stain (10,000x; Thermo Fisher Scientific, Merelbeke-Melle, Belgium) dissolved in 0.2 µm filtered dimethyl sulfoxide (Merck, Darmstadt, Germany). The viability staining mix (1.5) contained SYBR® Green I (10,000x) and Propidium Iodide (20 mM; Thermo Fischer Scientific, Merelbeke-Melle, Belgium) dissolved in 0.2 µm filtered dimethyl sulfoxide (Merck, Darmstadt, Germany). Samples (25 µL) were measured with the Fast fluidics setting on a volumetric BD Accuri^TM^ C6 Plus flow cytometer (BD Biosciences, Erembodegem, Belgium) with an event rate between 200 and 2000 events s^-1^, equipped with four fluorescence detectors (530/30 nm, 585/40 nm, >670 nm and 675/25 nm), forward and side scatter detectors, a 20 mW 488 nm laser and a 640 nm laser. A threshold of 500 was set to record signal on the green fluorescent channel (A).

Mock community with (1.4) or without (1.3) spiked DNA, human feces, saliva, the Simulator of the Human Intestinal Microbial Ecosystem (SHIME) gut reactor, soil and water samples (1.3, 1.4) and fresh and stored human and murine fecal suspensions (1.5) were sorted with a BD FACSMelody^TM^ cell sorter (BD Biosciences, Erembodegem, Belgium), equipped with six fluorescence detectors (527/32 nm, 700/54 nm, 582/15 nm, 613/18 nm, 697/58 nm, and 783/56 nm), one scatter detector (488/15 nm) and two lasers (488 nm and 561 nm). The sample line was disinfected after daily use with 70 % ethanol and a deep clean with bleach was performed weekly. The sheath fluid 0.2 µm inlet filter (BD Biosciences, Erembodegem, Belgium) was replaced regularly. A 100 µm sorting nozzle was used and sorting was performed in ‘Purity’ precision mode to exclusively select cell-containing droplets. All samples were diluted in filtered aPBS to an appropriate range for the cell sorter (10^5^ – 10^6^ cells mL^-1^) and stained with SYBR® Green I nucleic acid stain as described before. The event rate ranged between 200 and 6,000 events s^-1^, the sorting efficiency was above 80 %, the drop frequency was 34.0 kHz and a threshold was set around 100 on the green fluorescent channel. SYBR® Green I-stained samples were sorted into a SYBR® Green I-positive cell fraction and a cell-free noise fraction based on the gating of blue laser (detector A vs B) biplots (Figure S9) in the BD FACSChorus^TM^ software version 1.1.20.0. SYBR® Green I– and Propidium Iodide-stained samples were sorted into a SYBR® Green I-positive intact cell fraction and a double stained SYBR® Green I– and Propidium Iodide-positive damaged cell fraction based on the gating of blue laser biplots (Figure S10) in the BD FACSChorus^TM^ software version 1.1.20. Heat-killed samples (30 minutes at 96 °C), aPBS and 0.2 μm-filtered (Merck, Darmstadt, Germany) samples were included to inform the gating process that distinguished intact from damaged microbial cell subpopulations and cell-free noise. The sorted cell fraction consisted of 288,000 cells collected in 1 mL FACS sheath fluid in 1.5 mL Eppendorf® DNA LoBind collection tubes (Merck, Darmstadt, Germany). Data was further analyzed in R version 4.5.0 (3.2).

#### 2.4 PacBio full-length 16s rRNA gene sequencing

Prior to PacBio full-length 16s rRNA gene sequencing, DNA concentrations were quantified using a Qubit Flex fluorometer (Invitrogen) using the Qubit™ 1X dsDNA High Sensitivity Assay Kit by VIB Nucleomics Core (Leuven, Belgium). The 16S rRNA gene was amplified with the forward (27F 5’AGRGTTYGATYMTGGCTCAG) and reverse (1492R 5’RGYTACCTTGTTACGACTT) primer pair and converted to a Kinnex SMRTbell according to protocol PN 103-238-800 REV02 MAR2024. The SMRTbell was sequenced on a PacBio Sequel IIe device using the Sequel II binding kit 3.2. Data was further analyzed in R version 4.5.0 (3.3).

#### 2.5 Illumina 16s rRNA gene sequencing

Genomic DNA extract (10 µL) was sent to LGC genomics GmbH (Berlin, Germany) for sequencing and amplification of the 16S rRNA gene V3-V4 hypervariable region. The PCR mix included 1 µL of high or 6 µL of low microbial biomass DNA extract, 15 pmol of both the forward (341F 5’-CCTACGGGNGGCWGCAG) and reverse (785R 5’-ACTACHVGGGTATCTAAKCC) primer (Klindworth et al., 2013) in 20 µL MyTaq^TM^ buffer containing 1.5 U MyTaq^TM^ DNA polymerase (Bioline, Londen, UK) and 2 µL of BioStabII PCR Enhancer (Merck, Darmstadt, Germany). For each sample, the forward and reverse primers had the same unique 10-nt barcode sequence. PCR settings for high biomass DNA extracts were initial denaturation for 2 minutes at 96 °C, followed by 30 cycles of 15 seconds denaturation at 96 °C, annealing for 30 seconds at 50 °C, and extension for 90 seconds at 70 °C. Low biomass DNA was PCR amplified for 33 cycles. Agarose gel electrophoresis confirmed the sufficiently high yield of the amplicons of interest. Amplicon pools with 20 ng amplicon DNA of each sample for up to 48 samples carrying different barcodes were purified with one volume AMPure XP beads (Beckman Coulter Life Sciences, Brea, CA, USA) to remove primer dimers and other mispriming products, followed by an additional purification on MinElute columns (Qiagen, Venlo, The Netherlands). Illumina libraries consisting of 100 ng of each purified amplicon pool were constructed by means of adaptor ligation using the Ovation Rapid DR Multiplex System 1-96 (NuGEN) and size-selected by preparative gel electrophoresis. Sequencing was performed on an Illumina MiSeq instrument using v3 Chemistry (Illumina, San Diego, CA, USA). Data was further analyzed in R version 4.5.0 (3.3).

#### 2.6 Sanger 16S rRNA gene sequencing

The 16S rRNA gene from *Veillonella atypica* (Rogosa 1965) was Sanger sequenced to serve as a reference for the metataxonomic identification of *V. atypica* relic DNA contamination during FACS (1.4). Prior to Sanger sequencing, the 16S rRNA gene was amplified by PCR with the forward (27F 5’-GAGTTTGATCMTGGCTCAG) and reverse (1492R 5’-GGYTACCTTGTTACGACTT) primer pair (Eurogentec, Serraing, Belgium) in a T100^TM^ Thermal Cycler (Bio-Rad Laboratories Inc., Hercules, CA, USA). The reaction contained 1.25 µL of each primer (0.5 µM final concentration), 12.5 µL 2X Phusion Plus PCR Master Mix (Thermo Fisher Scientific, Merelbeke-Melle, Belgium), 9 µL Molecular Biology Grade Water (Labconsult, Ghent, Belgium) and 1 µL DNA extract. PCR amplification consisted of a pre-denaturation step of 30 seconds at 98 °C, followed by repeated denaturation (0 seconds at 98 °C), annealing (10 seconds at 55 °C), and extension (1 minute at 72 °C) for 30 cycles, followed by 5 minutes at 72 °C. PCR products were purified with the GeneJet PCR Purification Kit (Thermo Fisher Scientific, Merelbeke-Melle, Belgium). Purity of the PCR product was verified on a 2 % agarose gel (electrophoresis for 30 minutes at 100 V). The purified amplified DNA was sent to LGC Genomics GmbH (Berlin, Germany) for bi-directional Sanger sequencing. The quality of the forward and reverse 16S rRNA gene Sanger sequence chromatograms was inspected with BioEdit (Hall, 1999). The forward and reverse sequences were classified with the Ribosomal Database Project Naïve Bayesian Classifier (Cole et al., 2014; Quast et al., 2013) and aligned against 16S rRNA gene sequences using the BLAST algorithm for highly similar sequences (NCBI, accession date: June 2023) (Altschul et al., 1990; Johnson et al., 2008). Data was further analyzed in R version 4.5.0 (3.3).

### 3 Data analysis

We used R version 4.5.0 for data processing, visualization, and statistical analysis in RStudio 2025.05.0.

#### 3.1 qPCR

qPCR data (Quantstudio^TM^ 3; 2.1) was imported into RStudio. ROX and SYBR green fluorescent signals were stable. Amplification and melt curves were visualized in samples, standards and blanks on a linear and logarithmic scale using *ggplot2* (version 3.5.2, Wickham, 2011) to exclude samples with technical artifacts and non-specific amplification. Standard curves were constructed with linear regression and used to determine the 16S rRNA gene copy number and the amplification efficiency.

#### 3.2 Flow cytometry cell counting and FACS

Flow cytometry and FACS data were processed using the *ggcyto* (Van et al., 2018), *flowCore* (Hahne et al., 2009) and *openCyto* (Finak et al., 2014) R packages (*Bioconductor*, version 3.21). A biexponential transformation was performed for the purpose of visualization with *ggplot2*. Manually inspected forward scatter, side scatter, and fluorescence plots for all samples and blanks indicated a minimal carry-over and an adequate signal to noise ratio. Gating of the subpopulations was conducted as stated before (2.3, Figure S9 and S10).

#### 3.3 Metataxonomic bioinformatics analysis

The *DADA2* R package (version 1.36.0) was used to process the Illumina V3-V4 amplicon and PacBio full-length 16S rRNA gene sequencing data as described by Callahan et al. (2016). In a first quality control step, primer sequences were removed and reads were truncated at a quality score cut-off truncQ=2. Besides trimming, additional filtering was performed to eliminate reads containing any ambiguous base calls or reads with high expected errors (maxEE=2,2). Reads mapping to the Phix genome were discarded. After dereplication, unique reads were denoised using the Divisive Amplicon Denoising Algorithm (DADA) error estimation algorithm and the sample inference algorithm with the option pool=FALSE and singleton=FALSE. Denoised reads were merged after approval of the error rates. Finally, the amplicon sequencing variant (ASV) table obtained after chimera removal was used for taxonomy assignment using the Naive Bayesian Classifier (with an 80 % minimal bootstrap confidence threshold) and the DADA2 formatted Silva v138.1 database (Quast et al., 2013). For Illumina V3-V4 short-read sequencing data, species taxonomy assignment was based on exact matches to a species-specific database of Silva v138.1. For PacBio full-length 16S rRNA gene sequencing data, a minimal bootstrap of 80 % was used for species classification. The top 14 most abundant ASVs were selected using the *phyloseq* R package (version 1.52.0) (McMurdie & Holmes, 2013) and visualized at species and genus level using *ggplot2*. Further data analysis is elaborated in section 3.3 and 3.4.

Some specific DADA2 ASV sequences were aligned against the default Nucleotide collection using the NCBI Standard Nucleotide Basic Local Alignment Search tool (BLAST, version 2.15.0) with the megablast option, using default settings (Altschul et al., 1990; Johnson et al., 2008).

#### 3.4 *In silico* decontamination of FACS-sequencing data

*In silico* decontamination of the FACS-sequencing data was performed based on contaminating ASVs present in the sheath fluid and aPBS blanks. The identified ASVs in the blanks with an abundance below 0.1 % in high biomass references and mock samples were removed. This threshold prevents removal of carry-over ASVs that are abundant in ecosystem samples.

#### 3.5 Metataxonomics multivariate analysis

Metataxonomic sequencing data of diluted or sorted low biomass samples and high biomass references (1.2, 1.3, 1.4, 1.5) was compared by calculating the Jaccard absence/presence and abundance-weighted microbial community similarity using the *vegan* R package (version 2.6.10) (Jaccard, 1912; Oksanen et al., 2025). The obtained similarity matrices were visualized with Principal Coordinates Analysis (PCoA) using the *vegan* package (version 2.6-10). Observed Chao species richness was estimated using the *iNEXT* package (version 3.0.1) (Chao, 1984). *DESeq2* (version 1.48.1) was applied on the decontaminated unnormalized FACS-sequencing data of the stored fecal samples (1.5) to detect statistically significant differences in ASV abundances between the sorted fractions of the fresh and the stored slurry or feces (Love et al., 2014; McMurdie & Holmes, 2014). An empirical Bayes shrinkage correction was employed for low counts (Love et al., 2014). Pairwise significant differences were obtained using Wald tests, contrasting the sorted fresh feces to the sorted stored feces with and without glycerol-TE cryoprotectant. Significant differences were visualized in a volcano plot, showing the –log10(adjusted p-value) as a function of the shrunken log2 Fold Change. Genera or species with an adjusted p-value smaller than 0.05, were annotated in the plot (Quackenbush, 2002).

### 4 Data availability

Sequencing data have been deposited in the European Bioinformatics Institute’s (EBI) European Nucleotide Archive (ENA) with the accession code PRJEB90999. Flow cytometry data can be accessed on Zenodo with doi: 10.5281/zenodo.15747669.

## Results

We optimized and validated a FACS-sequencing workflow to remove contamination and bias in metataxonomic data from sorted microbial subpopulations with a low biomass.

### DNA yield of the PowerSoil® Pro kit increases with an increased bead beating frequency and sample input volume

The DNA extraction procedure was finetuned to maximize DNA yields from sorted low microbial biomass samples. The DNeasy® PowerSoil® Pro kit was selected as a basis to optimize DNA extraction of low biomass samples, since it outperforms other methods such as Chelex or mechanical lysis followed by phenol/isopropanol/ethanol purification (Demkina et al. 2023). The PowerSoil® Pro manufacturer’s protocol was modified by testing an increased sample input volume from 250 to 500 µL in combination with an increased bead beating frequency from 2,000 rpm in two intervals of 30 seconds with 30 seconds cooldown to 4,000 rpm for five consecutive intervals of 15 seconds with 45 seconds cooldown. The combined effect of an increased 500 µL sample input volume and 4,000 rpm bead beating frequency resulted in the highest 16S rRNA gene copy number in DNA extracts from *Bacillus subtilis* (147.3*10^3^ ± 9.9*10^3^ 16S rRNA gene copies mL^-1^) and *Escherichia coli* (38.0*10^3^ ± 0.5*10^3^ 16S rRNA gene copies mL^-1^) sorted cells (Figure 1). This represented a 2.65-fold increase compared to the PowerSoil Pro reference protocol using 250 µL and 2,000 rpm. Input volume was the biggest driver of the combined effect. The protocol using 500 µL at 2,000 rpm performed second best resulting in a 1.62-fold increase compared to the reference. The 250 µL 4,000 rpm modification negatively affected the DNA extraction yield compared to the reference, decreasing the copy number 12.7-fold. Non-sorted high and diluted low biomass *B. subtilis* and *E. coli* samples and sorted noise showed consistent results, confirming the applicability of the modified PowerSoil Pro DNA extraction protocol to extract DNA from a wide range of microbial cell concentrations. Comparable results were obtained using less precise fluorometric dsDNA quantification (Figure S1).

**Figure 1:**
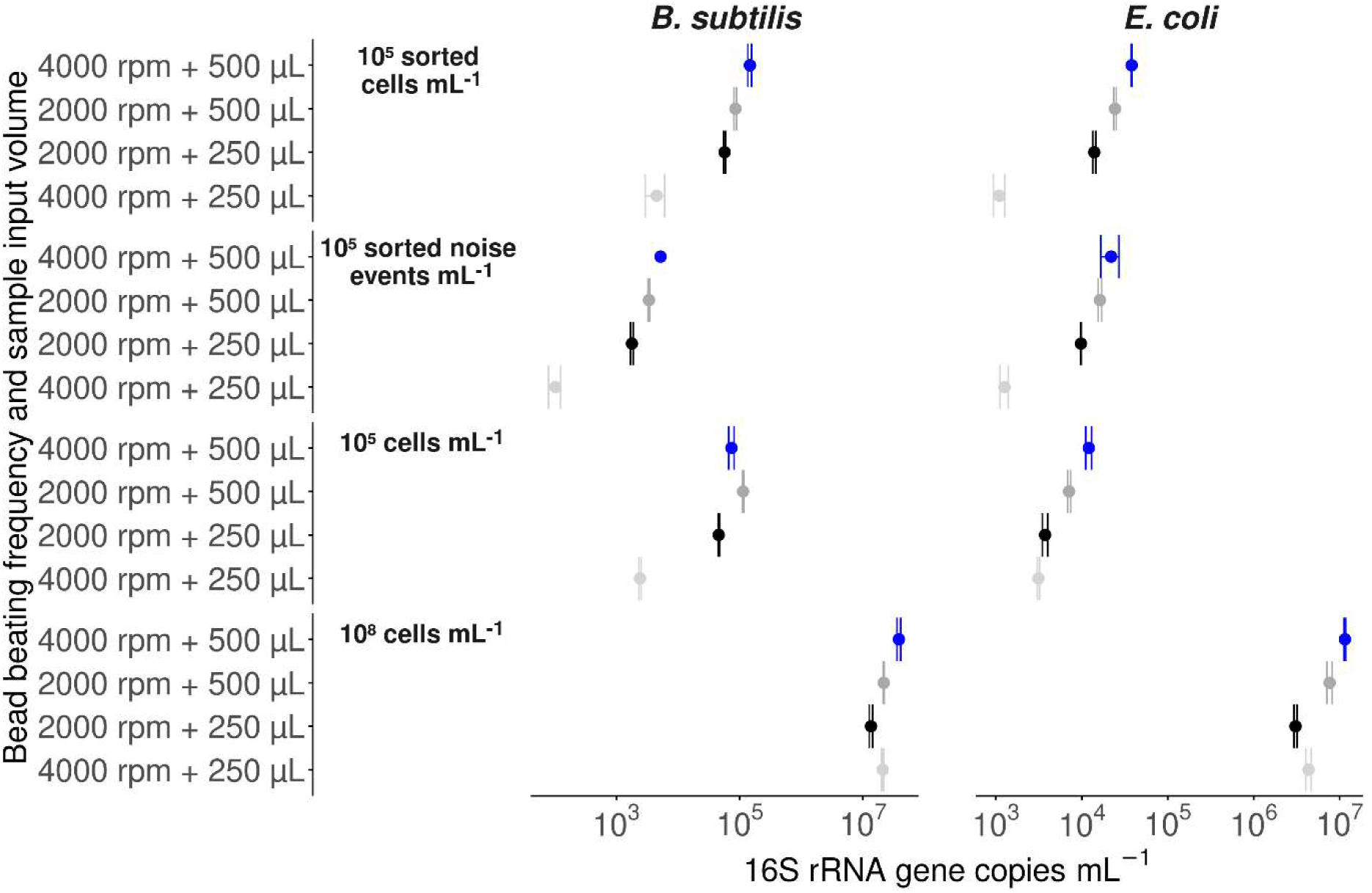
The PowerSoil® Pro protocol with an increased 500 µL input volume and 4,000 rpm bead beating (blue) yielded the highest average (n=3) qPCR 16S rRNA gene copy number in sorted (∼ 2.88 * 10^5^ cells mL^-1^), diluted (∼ 10^5^ cells mL^-1^) and high biomass reference (∼ 10^8^ cells mL^-1^) samples and sorted noise (∼ 2.88 * 10^5^ noise events mL^-1^) derived from pure cultures of *Bacillus subtilis* subsp. *subtilis* strain 168 and of *Escherichia coli* str. K-12 substr. MG1655. The protocol using 500 µL at 2,000 rpm (dark gray) performed second best. The 250 µL 4,000 rpm modification (light grey) negatively affected the DNA extraction yield compared to the reference (black).

The superior PowerSoil Pro DNA extraction protocol with 500 µL sample input volume and 4,000 rpm bead beating was further evaluated by comparing the microbial community composition in 10^5^ cells mL^-1^ diluted low biomass and 10^7^ cells mL^-1^ high biomass reference samples from feces, saliva, the Simulator of the Human Intestinal Microbial Ecosystem (SHIME) gut reactor, soil and water microbial ecosystems.

### The PowerSoil® Pro kit protocol with an increased bead beating frequency and sample input volume effectively extracts DNA from low biomass samples obtained from feces, saliva, the SHIME gut reactor, soil and water microbial ecosystems

The optimized PowerSoil Pro DNA extraction protocol with 500 µL sample input volume and 4,000 rpm bead beating successfully extracted DNA from diverse high biomass (10^7^ cells mL^-1^) and diluted low biomass (10^5^ cells mL^-1^) microbial ecosystems. PacBio full-length 16S rRNA gene sequencing of the low biomass DNA extracts yielded higher read counts compared to blank samples of the anaerobic PBS (aPBS) diluent but reduced read counts compared to the high biomass reference DNA extracts obtained from saliva, the Simulator of the Human Intestinal Microbial Ecosystem (SHIME) gut reactor, feces, soil and water microbial ecosystems (Table S1).

The lower read counts corresponded to a reduction in observed Chao species richness (Table S2) with 7 (saliva), 13 (SHIME), 58 (feces), 388 (soil) and 365 (water). The richness reduction was larger in soil and water samples with a high richness of 471 and 423 species, respectively, in the high biomass reference samples. Feces, saliva and SHIME samples with a lower richness of 109, 121 and 47 species, respectively, displayed a smaller decrease in richness due to biomass dilution. This resulted in higher Jaccard similarity in species absence/presence between high and low biomass samples from saliva (67.9 %), SHIME (55.8 %) and feces (33.3 %) compared to soil (15.9 %) and water (10.6 %) microbial ecosystems. While biomass dilution largely affected the detection of rare taxa and absence/presence based community similarity, it had less effects on the abundant taxa (Figure S2). The abundance-weighted Jaccard similarity reached 66.4 % in saliva, 72.7 % in SHIME, 52.2 % in feces, 38 % in soil and 51 % in water.

Illumina V3-V4 amplicon 16S rRNA gene sequencing of low biomass samples likewise yielded a lower absence/presence-based versus abundance-weighted microbial community similarity (Figure S3). In contrast to PacBio full-length 16S rRNA gene sequencing, the Chao species richness increased upon microbial biomass dilution from 98 to 123 in saliva, from 78 to 249 in soil and from 57 to 242 in water (Table S2). The increased richness resulted from an increased detection of rare taxa and coincided with increased read counts in low microbial biomass saliva (137,570 versus 54,681) and water (21,742 versus 8,941) samples (Table S1). The increased read counts were likely due to an increased amplicon product yield resulting from increased DNA input (6 versus 1 µL) in combination with three additional PCR cycles for low biomass samples. The DNA input used for PacBio sequencing was generally higher in high biomass samples. The amplicon product yield is also expected to be higher in high biomass samples since amplification was limited to 20 PCR cycles in both high and low biomass samples (Table S1). The richness thus depended on sequencing read depth which, in turn, depended on the amplicon product yield obtained during sequencing library preparation. Carry-over from co-analyzed high biomass samples from other microbial ecosystems did not contribute to the increased richness since the rare taxa detected in the Illumina sequenced low biomass saliva, soil and water samples were not detected in any of the sequenced high biomass samples.

To rule out that the increased richness in the Illumina sequenced diluted low microbial biomass samples was due to human-derived or laboratory contamination, the validated protocol was further used to extract control samples. These controls were also used to detect and remove contamination in the sequencing data of the sorted low biomass samples from the various microbial ecosystems in order to improve the accuracy of FACS-sequencing.

### FACS-sequencing introduces contaminants in the sequencing data of the sorted low biomass microbiota

Contamination detection in sorted low microbial biomass samples relied on PacBio and Illumina 16S rRNA gene sequencing of non-selectively sorted ZymoBIOMICS mock community samples consisting of a mixture of eight inactivated bacterial species, FACS sheath fluid used to create a single-cell suspension in the sorter and aPBS used as diluent.

FACS-sequencing introduced a set of contaminants, further termed the FACSome, in sorted microbiota. Sorted mock microbiota were consistently contaminated with three FACSome ASVs that were traced back to FACS sheath fluid: *Alcaligenes faecalis* ASV118, *Aquamicrobium* sp. ASV 73 and *Pseudomonas veronii,* ASV39 (Figure 2). *P. veronii* was also consistently observed to contaminate sorted low biomass microbiota from feces, saliva, the Simulator of the Human Intestinal Microbial Ecosystem (SHIME) gut reactor, soil and water ecosystems (Figure S2). *A. faecalis* and *Aquamicrobium* sp. occurred in 6 and 4 out of the 9 sorted ecosystem samples, respectively. The FACSome ASVs were exclusive to sorted samples and were detected in both the sorted cell and cell-free noise fractions, suggesting that this contamination at least partly originated from extracellular relic DNA. Other contaminating ASVs were not exclusive to the sorted samples.

**Figure 2:**
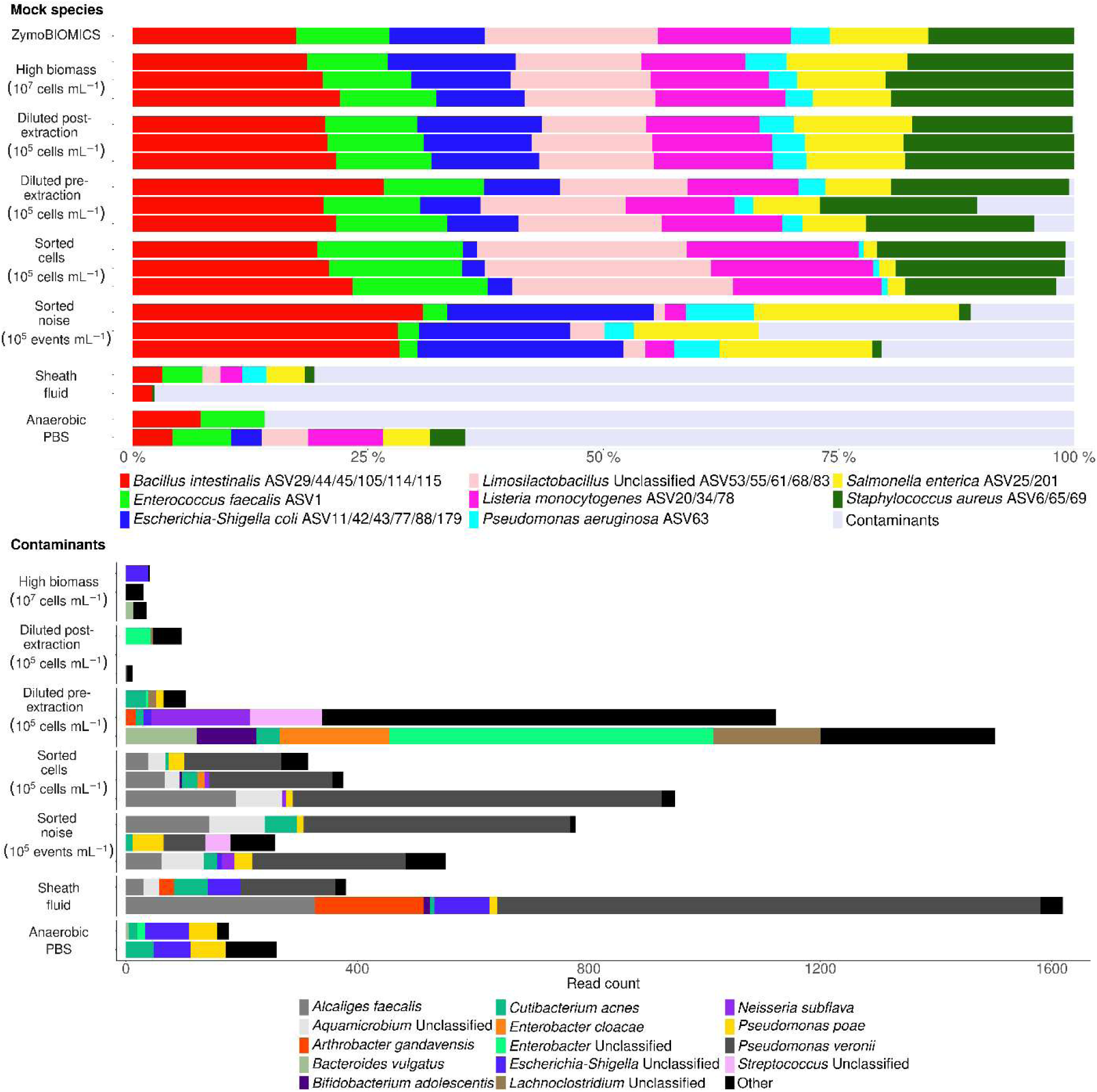
Contaminants were detected in the metataxonomic sequencing data of low biomass ZymoBIOMICS mock microbiota. Samples were sequenced with PacBio full-length 16S rRNA gene sequencing. The low biomass was obtained through dilution of the high biomass mock standard (10^7^ cells mL^-1^) to 10^5^ cells mL^-1^ before (diluted pre-extraction) and after (diluted post-extraction) DNA extraction and through sorting of the mock standard into 10^5^ cells mL^-1^ and 10^5^ noise events mL^-1^ using a non-selective SYBR® Green I staining. Dilution and sorting were performed in triplicate. FACS sheath fluid and aPBS diluent blanks were included in duplicate to identify and track the FACSome contaminants introduced during FACS-sequencing.

The human skin commensal *Cutibacterium acnes* ASV1242/9362 was detected in 5 out of the 6 sorted mock samples, 4 out of the 9 sorted ecosystem samples, one sheath fluid blank and both aPBS blanks. It was absent in high biomass mock samples (n=3) and DNA extracts from high biomass mock samples that were diluted to concentrations corresponding to 10^5^ cells mL^-1^ after extraction (n=3). Dilution (n=3) of mock samples to 10^5^ cells mL^-1^ before extraction introduced *C. acnes*. The same trend was observed for the plant-associated *Pseudomonas poae* ASV1262/2809/3664/6421. While both taxa were not exclusive to sorted samples they were introduced by FACS-sequencing since FACS generated low microbial biomass samples that were susceptible to *P. poae* and *C. acnes* contamination that was detected through sequencing.

Sorted low microbial biomass samples were also prone to contamination by human salivary and fecal bacteria. The salivary bacterium *Neisseria subflava* (ASV7881/8307) was detected in half of the sorted mock samples and one noise sample contained salivary *Streptococcus* sp. (ASV4566). A single sorted mock sample contained fecal *Enterobacter cloacae* (ASV16268). Different ASVs of these salivary (*Streptococcus* sp. ASV4/638/798/1205/1223/1823/3311/12333/14594 and *N. subflava* ASV134/4474) and fecal (*E. cloacae* ASV139) species were retrieved in some of the samples that were diluted to a low biomass (10^5^ cells mL^-1^) prior to extraction. The ASVs in these diluted samples matched the ASVs detected in saliva (*Streptococcus* ASV4/638/798/1205/1223/1823/3311 and *N. subflava* ASV134/4474), respectively, SHIME gut reactor samples (*E. cloacae* ASV139) that were processed in the same experimental run. A closer inspection of the pre-extraction diluted low biomass samples revealed that one sample shared 33 ASVs with the saliva sample and another shared 22 ASVS with the SHIME gut reactor sample (Figure S2), suggesting that these contaminants were introduced by carry-over of sample or extracted DNA. Carry-over was not evidenced for the salivary and fecal ASVs that contaminated the sorted samples, which were extracted separated in time and space from the high biomass references.

Carry-over was, furthermore, reduced in the samples diluted to a DNA concentration corresponding to 10^5^ cells mL^-1^ after extraction (36 ± 53 carry-over reads) compared to samples diluted to 10^5^ cells mL^-1^ before extraction (910 ± 723 carry-over reads) and was absent in the 10^7^ cells mL^-1^ high biomass mock samples. This demonstrates that low biomass samples are susceptible to carry-over and that manipulation of the low biomass samples or DNA extracts in the presence of high biomass samples permitted the carry-over. Carry-over inflated the percentage of contaminating reads. The pre-extraction diluted samples affected by carry-over contained 1,502 and 1,123 contaminating reads, while 436 ± 361 (n=4) contaminating reads were found in another replicate (n=1) and all sorted mock microbiota replicates (n=3) that did not display carry-over (Figure 2). All of the reads detected in the sheath fluid (1658 and 471) and aPBS (402 and 207) blanks were contaminating and 10.8 % ± 12.0 % (n=2) and 24.7 % ± 15.1 % (n=2) of the reads resulted from mock carry-over, respectively.

Carry-over in sheath fluid and aPBS samples mostly consisted of mock ASVs. Besides, the SHIME-derived *Escherichia*-*Shigella* sp. ASV101 was detected in sheath fluid and aPBS.

*Escherichia*-*Shigella sp*. ASV1272 co-existed in these blanks, as well as, sorted samples but did not originate from the SHIME gut reactor samples. Instead, this ASV mapped onto the genome of *Escherichia coli* str. K-12 substrain MG1655 with 99.8 % identity and a BLAST® E-value of 0. *E. coli* str. K-12 substrain MG1655 is a common lab strain, which was also used to optimize DNA extraction in the first experiment that was performed in the same lab environment weeks before the sorting experiment took place. This illustrates that sorted low biomass samples are vulnerable to the acquisition of contaminating cells and DNA from the lab environment.

*Arthrobacter gandavensis* ASV1782, found in both sheath fluid replicates, one fecal sample and one mock community sample diluted before DNA extraction also qualifies as a lab contaminant. The ASV corresponded with a cultured isolate (BLAST® 99.9 % sequence identity, E-value of 0) that was processed in the DNA extraction lab. It is not part of the FACSome as it was not detected in any of the sorted samples.

Other contaminating ASVs that were not incorporated in the FACSome and displayed a low abundance in some of the blank and diluted samples were summarized in Table S3-S5.

The same samples were also sequenced on an Illumina sequencing platform using V3-V4 primers (Figures S3-S6 and Tables S6-S8). Similar data, albeit with lower read counts and lower taxonomic resolution, was obtained compared to PacBio sequencing. The identified FACSome contaminant sequences were identical to and perfectly aligned with PacBio full-length 16S rRNA gene sequences with 100 % BLAST identity. Illumina sequencing data was enriched in *Cutibacterium* reads, while carry-over contaminants were absent in the samples diluted to 10^5^ cells mL^-1^ prior to extraction.

While sample carry-over can be minimized by banning high biomass samples from the workspace when handling low biomass samples and DNA extracts, contaminations introduced during the sorting procedure or the human manipulation of samples in the lab environment are difficult to prevent in standard lab facilities. We, therefore, developed an *in silico* data decontamination strategy guided by the identification of contaminants in mock microbiota, sheath fluid and aPBS blanks to increase the accuracy of FACS-sequencing.

### *In silico* decontamination of FACS-sequencing data increases accuracy

The removal of contaminant ASVs increased the accuracy of the microbial community composition determined with PacBio and Illumina 16S rRNA gene sequencing in sorted low biomass samples.

*In silico* decontamination based on sheath fluid and aPBS blanks eliminated the FACSome and other human-derived and laboratory contaminants while preserving the mock species in sorted ZymoBIOMCS mock microbiota (Figure 3). The removal of contaminant ASVs increased the Jaccard similarity between the sorted and the theoretical mock microbiota composition (Figure 4 and Figure S7) from 47.2 % ± 2.8 % to 62.8 % ± 4.8 % (n=3).

**Figure 3:**
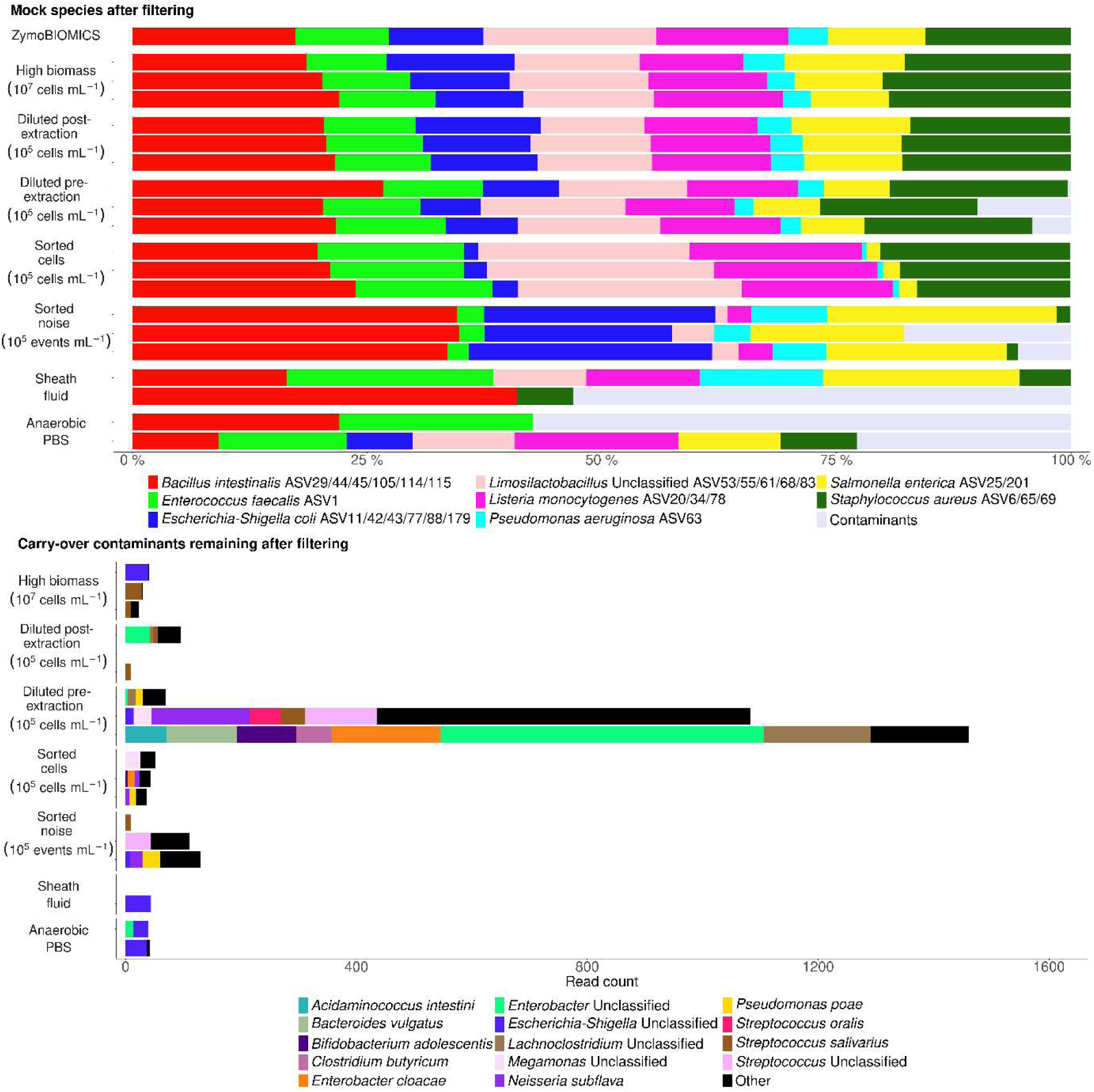
*In silico* data decontamination removed contaminants from the metataxonomic sequencing data of low biomass ZymoBIOMICS mock microbiota. The removal of relic DNA during sorting resulted in discrepancies between the non-sorted and sorted samples. Samples were sequenced with PacBio full-length 16S rRNA gene sequencing. The low biomass was obtained through dilution of the high biomass mock standard (10^7^ cells mL^-1^) to 10^5^ cells mL^-1^ before (diluted pre-extraction) and after (diluted post-extraction) DNA extraction and through sorting of the mock standard into 10^5^ cells mL^-1^ and 10^5^ noise events mL^-1^ using a non-selective SYBR® Green I staining. Dilution and sorting were performed in triplicate. FACS sheath fluid and aPBS diluent blanks were included in duplicate to identify and track the FACSome contaminants introduced during FACS-sequencing.

**Figure 4:**
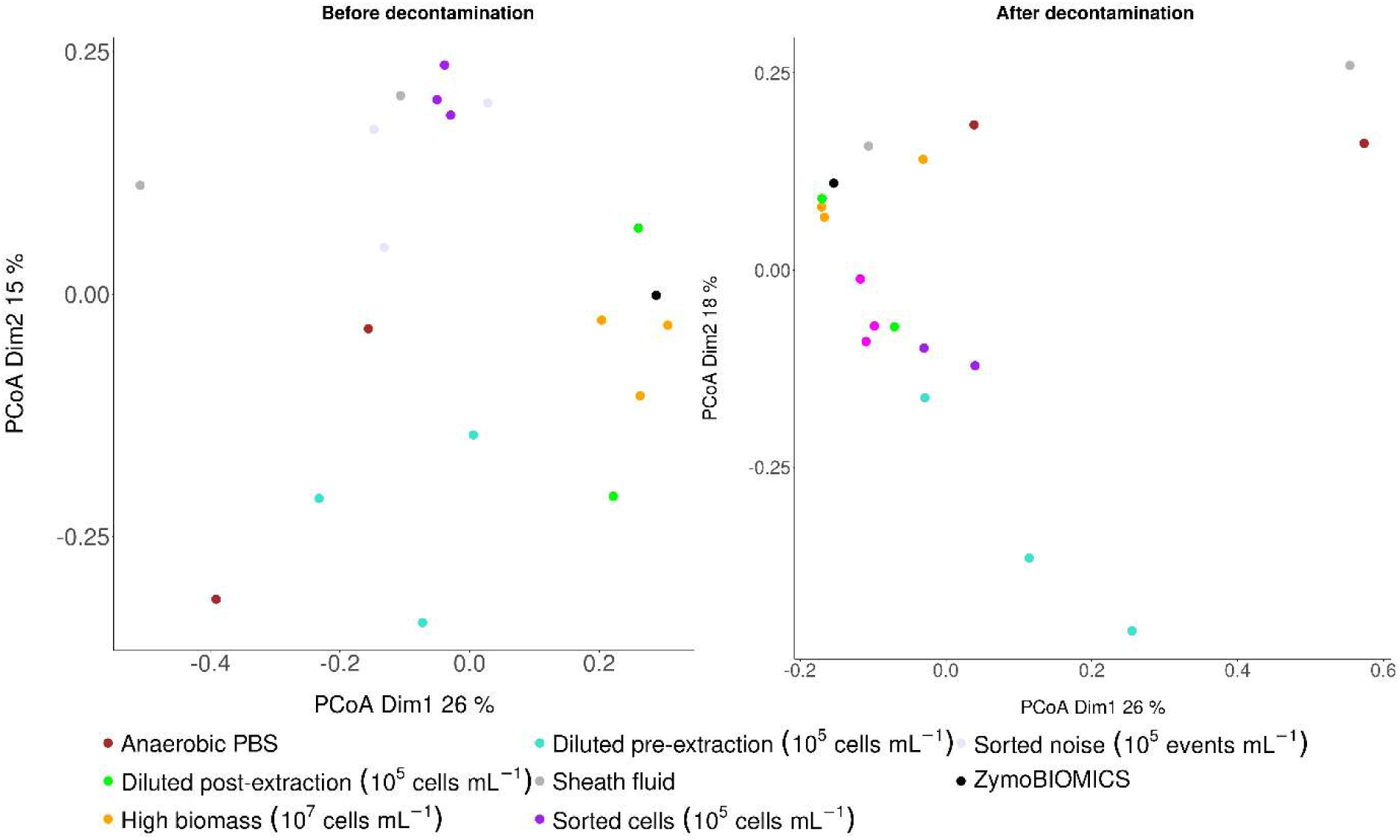
*In silico* data decontamination increased the absence/presence-based Jaccard similarity between sorted ZymoBIOMICS mock cells (10^5^ cells mL^-1^; pink) and the theoretical ZymoBIOMICS standard (black) resulting in a closer ordination of these samples in a Principal Coordinates Analysis (PCoA). Low biomass samples obtained through dilution of the high biomass mock standard (orange; 10^7^ cells mL^-1^) to 10^5^ cells mL^-1^ before (diluted pre-extraction; turquoise) and after (diluted post-extraction; green) DNA extraction and sorted noise events (10^5^ mL^-1^; purple) did not display an improved clustering. Samples were sequenced with PacBio full-length 16S rRNA gene sequencing. Dilution and sorting with a non-selective SYBR® Green I cell staining were performed in triplicate. FACS sheath fluid (light grey) and aPBS (brown) diluent blanks were included in duplicate to identify and track the FACSome contaminants introduced during FACS-sequencing.

The similarity of all post– and pre-extraction diluted mock samples (n=9) to the mock standard increased slightly from 61.1 % ± 27.5 % to 63.3 % ± 28.1 % following decontamination. This follows from the fact that these samples were not sorted and thus not exposed to FACS sheath fluid contaminants. Instead, the diluted mock samples were contaminated by fecal and salivary sample carry-over from co-analyzed high biomass references. These fecal and salivary carry-over contaminants were not removed as they were absent in the aPBS and sheath fluid samples used to identify the FACSome. The aPBS and sheath fluid samples did show evidence of mock sample carry-over. This carry-over did not affect sorted samples. Carry-over contaminants were excluded from the FACSome, by limiting the FACSome to those aPBS and sheath fluid contaminants that did not exceed a 0.1 % abundance in high biomass references. This threshold ensured that all mock species were retained in the *in silico* decontaminated sorted mock samples.

After decontamination, the sorted mock microbiota was still distorted compared to the mock standard. Gram-negative *Pseudomonas aeruginosa* ASV63, *Escherichia*-*Shigella coli* ASV11/42/43/77/88/179 and *Salmonella enterica* ASV25/201 had a decreased abundance in the sorted mock cells, but were enriched in the sorted noise fraction. This reduced abundance of easy-to-lyse Gram-negative genera in the sorted cell fraction and the concurrent detection of their released relic DNA in the noise fraction is consistent with lysis of Gram-negative cells by the DNA/RNA Shield^TM^ solution in which the mock community standard was stored prior to FACS. The bias in Gram-negative mock taxa was solely due to cell lysis during storage. Gram-negative genera such as *Alistipes* were successfully sorted out in the cell fraction when fresh fecal samples were used (Figure 5 and S2).

**Figure 5:**
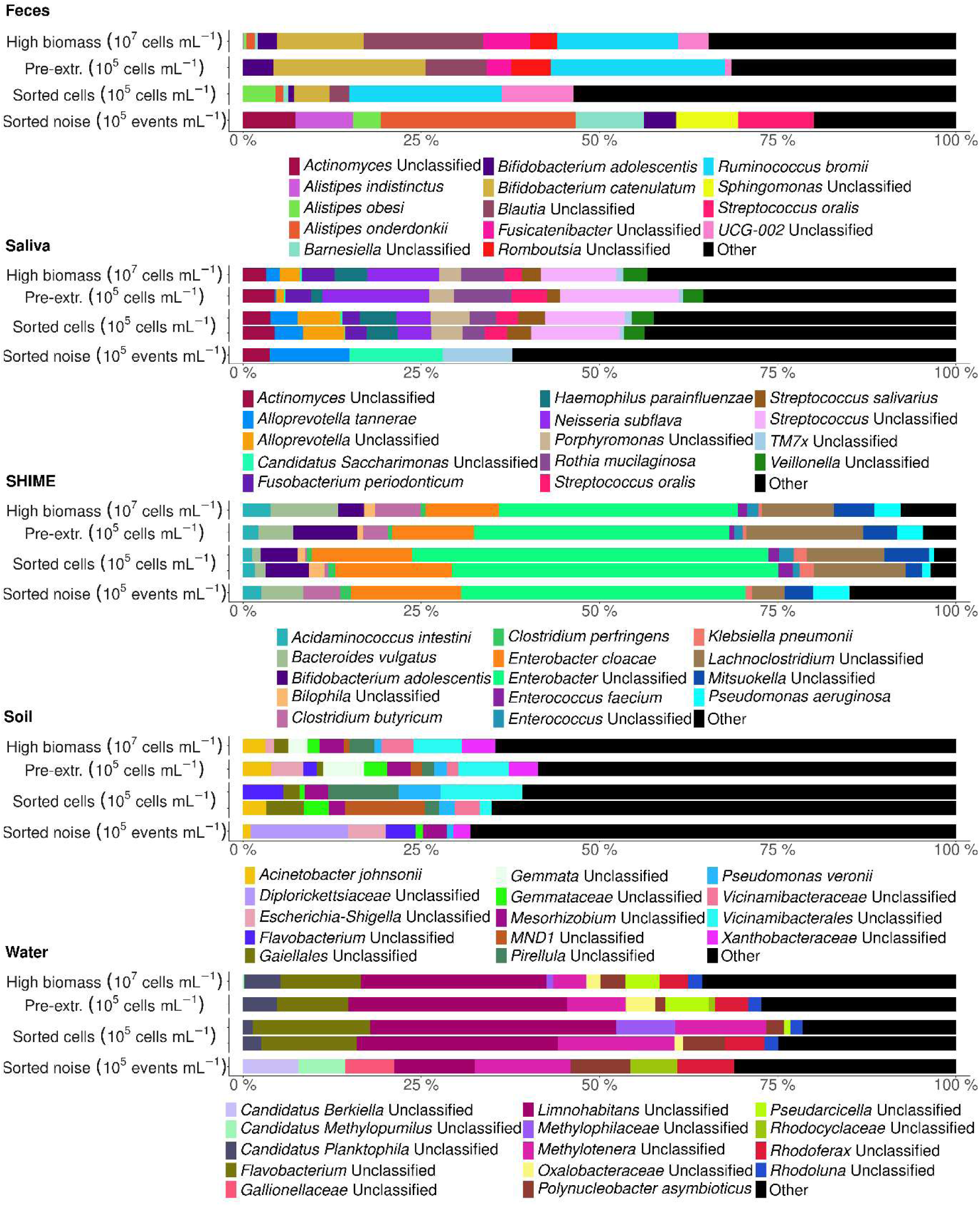
*In silico* data decontamination removed contaminants from the metataxonomic sequencing data of low biomass feces, saliva, SHIME gut reactor, soil and water microbiota. The removal of relic DNA during sorting resulted in discrepancies between the non-sorted and sorted samples. Samples were sequenced with PacBio full-length 16S rRNA gene sequencing. The low biomass was obtained through dilution of the high biomass reference samples (10^7^ cells mL^-1^) to 10^5^ cells mL^-1^ before extraction (Pre-extr.) and through sorting of the high biomass samples into 10^5^ cells mL^-1^ and 10^5^ noise events mL^-1^ using a non-selective SYBR® Green I staining. Dilution and sorting were performed in duplicate (except for feces).

Even fresh ecosystem samples can contain a large amount of relic DNA (Carini et al., 2016; Gryp et al., 2020), which was likely enriched in the noise and depleted or even eliminated in the cell fraction during FACS, causing discrepancies between the sequenced sorted cell fraction and the high biomass references and the samples diluted to 10^5^ cells mL^-1^ before DNA extraction. The elimination of relic DNA in the cell fraction caused a drop in observed Chao species richness of 45.5 (soil), 31.5 (water), and 8.5 (SHIME) compared to the diluted low biomass samples after decontamination. This drop was not due to subsampling since the diluted low biomass samples were diluted to 10^5^ cells mL^-1^ before extraction to mimic subsampling by sorting to 10^5^ cells mL^-1^. The reduced richness and additional depletion of relic DNA in the sorted cells resulted in the low similarity of the sorted soil (7.0 % ± 1.4 %; n=2) and water samples (4.9 % ± 1.3 %; n=2) compared to the high biomass references. Even after a successful *in silico* decontamination, there was no increase in similarity for soil and water samples (Figure 5 and S2). Similarity was higher in the feces (46.0 %; n=1), saliva (65.5 % ± 5.0 %; n=2) and SHIME (39.4 % ± 6.8 %; n=2) sorted communities, which exhibited a slight increase in similarity (47.5 %, 67.6 % ± 5.0 % and 41.7 % ± 7.3 %, respectively) after decontamination.

To confirm that the low similarities resulted from the depletion of relic DNA in the sorted cell fraction and did not reflect a low accuracy of FACS-sequencing compared to direct sequencing of diluted low biomass samples, a relic DNA spike-in experiment was performed.

### FACS filters out relic DNA contamination

Spike-in of a ZymoBIOMCS mock community with relic DNA from *Veillonella atypica* (Rogosa 1965) prior to FACS, confirmed the depletion of relic DNA in the sorted cell fraction and the concurrent emergence of relic DNA in the sorted noise fraction.

The relative abundance of *V. atypica* 16S sequences decreased from 9.3 %, corresponding to 3.5 ng spiked DNA in 10^7^ mock cells mL^-1^, to 0.3 % in the sorted mock cells (Table S9). *V. atypica* constituted 5.8 % of the sorted noise fraction, demonstrating the detection of relic DNA in the sorted noise (Table S9). Sorted noise was also enriched in easy-to-lyse Gram-negative *Pseudomonas aeruginosa* ASV63, *Escherichia*-*Shigella coli* ASV11/42/43/77/88/179 and *Salmonella enterica* ASV25/201 relic DNA, the FACSome contaminants *Alcaligenes faecalis* ASV118, *Aquamicrobium* sp. ASV73 and *Pseudomonas veronii* ASV39 and the human skin-derived contaminant *Cutibacterium acnes*, validating the reproducibility of our previous findings (Figure 6). Spiked PBS replicates did not contain cells but contained 47.7 and 17.0 % *V. atypica* in the noise fraction, along with FACSome contaminants and mock taxa resulting from carry-over. The FACSome contaminants included *A. faecalis* ASV118*, Aquamicrobium* sp. ASV73 *and P. veronii* ASV39 that were previously identified next to an unclassified *Bacillus* ASV787. This observed variation in the FACS sheath fluid composition illustrates the dynamic nature of the FACSome and substantiates the need for *in silico* decontamination data based on control samples that are co-analyzed in the same sorting run.

**Figure 6:**
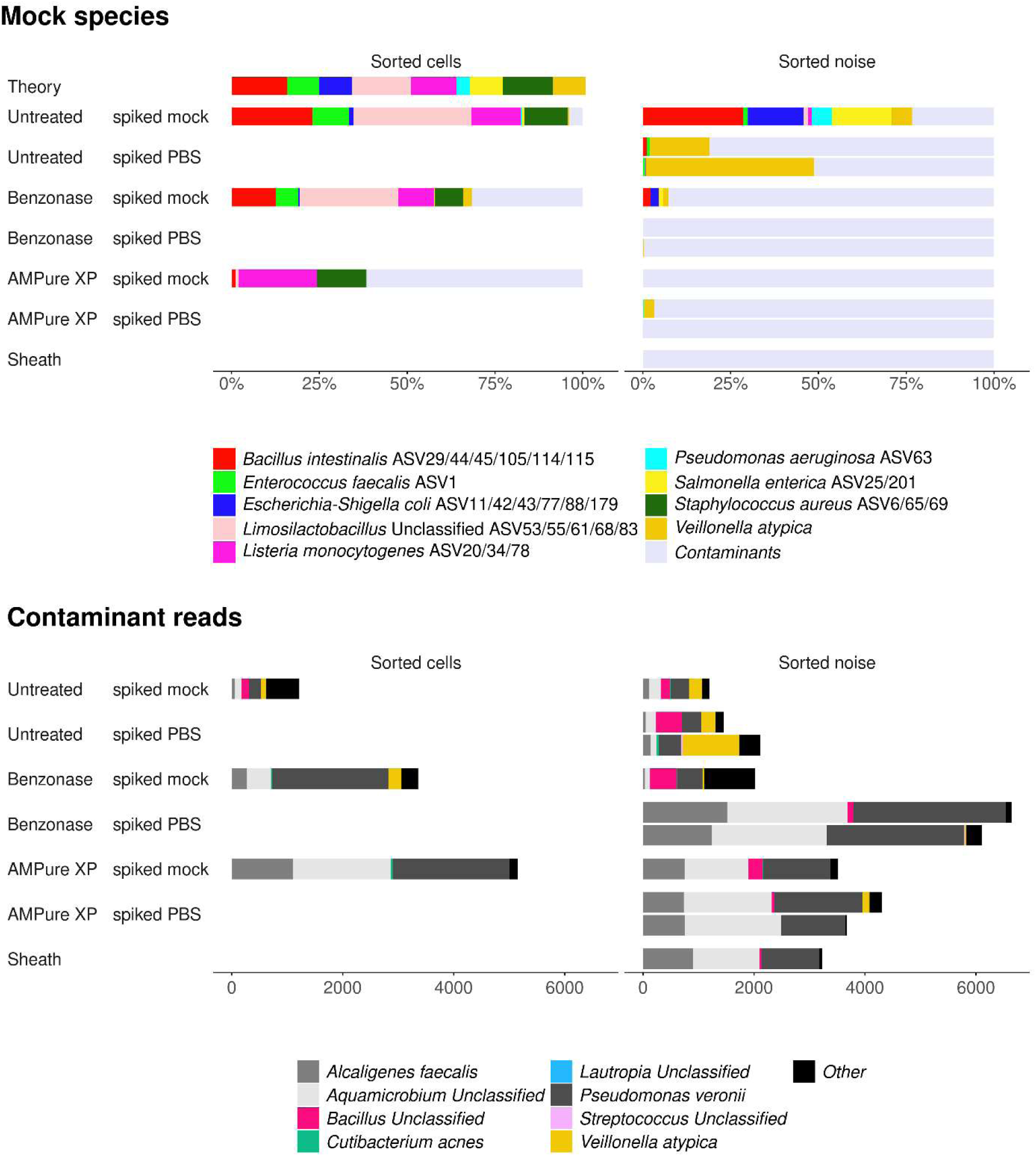
*Ve*illonella *atypica* (Rogosa 1965) relic DNA, spiked into a ZymoBIOMICS microbial community standard (mock) and PBS, was depleted in sorted microbial cells (10^5^ cells mL^-1^) and emerged in sorted noise (10^5^ noise events mL^-1^). Benzonase nuclease activity selectively depleted relic DNA of *Veillonella atypica*, *Pseudomonas aeruginosa* ASV63, *Escherichia*-*Shigella coli* ASV 11/42/43/77/88/179 and *Salmonella enterica* ASV 25/201. AMPure XP clean-up eliminated relic DNA but was not selective resulting in a distorted community composition. A decrease in the (relic) DNA concentration prior to sorting resulted in an inflation in absolute and relative contaminant sequencing read counts attesting the need of *in silico* data decontamination. Samples were sequenced with PacBio full-length 16S rRNA gene sequencing. Cells were sorted using a non-selective SYBR® Green I staining. A FACS sheath fluid blank was included to identify and track the FACSome contaminants introduced during FACS-sequencing.

Purification of samples with Benzonase prior to FACS removed 95.8 % of spiked *V. atypica* 16S sequences compared to 93.4 % removal through FACS in untreated sorted samples. The Benzonase nuclease activity additionally depleted *P. aeruginosa* ASV63, *E. coli* ASV11/42/43/77/88/179 and *S. enterica* ASV25/201 in the sorted cells and noise, which confirms that these easy-to-lyse Gram-negative species were present in the mock standard as relic DNA and not as intact cells. Spiked *V. atypica* DNA was reduced to 1.4 % in the noise fraction of the mock standard and to 0 and 0.3% in the spiked PBS replicates but reached 2.4 % in the mock cell fraction, indicating that the Benzonase incubation conditions need to be optimized to completely digest extracellular DNA in samples with high relic DNA concentrations. The removal of relic DNA in the spiked mock sample increased the relative importance of contaminants introduced downstream during FACS-sequencing which is reflected in a larger number and fraction of FACSome reads.

A similar inflation in absolute and relative contaminant read counts was observed in sorted cells and noise of mock and PBS samples treated with AMPure XP beads. AMPure XP beads eliminated spiked *V. atypica*, as well as, *P. aeruginosa* ASV63, *E. coli* ASV11/42/43/77/88/179 and *S. enterica* ASV25/201 relic DNA in sorted mock and PBS samples. The beads did not selectively remove relic DNA judged by the additional elimination of *Enterococcus* ASV1 and the near elimination of *Bacillus* ASV29/44/45/105/114/115 and *Limosilactobacillus* ASV53/55/61/68/83 compared to the Benzonase treated and untreated samples. The AMPure XP clean-up procedure thus distorted the microbial community composition and is not suitable for relic DNA removal.

Relic DNA filtering through FACS with additional Benzonase treatment, on the other hand, can improve the accuracy of function-driven microbial community analysis since relic DNA could interfere with targeted activity-, viability-, or function-targeted single-cell labelling and sorting. Activity– or viability-based microbial community profiling, furthermore, requires the preservation of intact viable cells during sample storage prior to FACS-sequencing. Storage effects on FACS-sequencing of the intact cell population were assessed for *in vivo*-derived murine and human fecal samples, which are routinely stored at –80 °C in microbiome cohort studies in which immediate analysis is hampered by practical constraints.

### Direct storage of fecal pellets reduces compositional shifts in sorted intact cells compared to storage of fecal slurries amended with cryoprotectants

Storage of *in vivo*-derived murine and human fecal samples at –80°C for one month reduced cell integrity and induced a compositional shift in the intact sorted *in silico* decontaminated microbiota compared to an immediately analyzed benchmark. Direct storage was superior to storage in 5 % glycerol – 1X Tris-EDTA (glycerol-TE) buffer, which has previously been used as a cryoprotectant in single-cell FACS-sequencing.

Direct storage of murine fecal pellets better preserved microbial viability compared to storage of murine fecal slurries with glycerol-TE. The intact cell ratio decreased from 74.8 % ± 0.1 % in fresh pellets to 65.4 % in stored pellets and 54.3 % ± 0.1 % in stored slurries. The absence/presence-based community similarity of the intact microbiota, calculated based on *in silico* decontaminated Illumina 16S rRNA gene sequencing data, was lowest (40.7 % ± 19.1 %; Figure S8) between stored murine fecal slurries and freshly analyzed benchmarks. Storage of murine fecal slurries in glycerol-TE significantly reduced *Lachnospiraceae* UCG-006 (Log Fold Change (LFC) = –5.7, p = 3.9 * 10^-2^)*, Acetatifactor* (LFC = – 5.3, p = 2.0 *10^-2^), *Akkermansia* (LFC=-5.2, p=1.7 * 10^-2^, *Prevotellaceae* UCG-001 (LFC = –6.0, p = 2.0 * 10^-2^) and *Faecalibacterium* sp. UBA1819 (LFC = –8.1, p = 1.7 * 10^-2^) abundance in the intact cell fraction (Figure 7). Direct storage had a smaller impact on the intact microbial community (Jaccard similarity of 53.7 % ± 0.7 %; Figure S8) and caused significant decreases in *Butyricicoccus* (LFC = –6.7, p = 4.9 * 10^-2^) and some *Lachnospiraceae,* A2 (LFC = –7.5, p = 9.0 * 10^-3^) and *Marvinbryantia* (LFC = –7.7, p = 9.0 * 10^-3^; Figure 7), next to a significant increase of *Eubacterium* (LFC = 8.8, p = 1.1 * 10^-4^) and *Candidatus Saccharimonas* (LFC = 7.1, p = 9 * 10^-3^). The duplicate murine intact microbial communities showed a high variability that was not introduced during storage or FACS since the variability was already present in the freshly analyzed benchmarks coming from two different fecal pellets (Figure 8).

**Figure 7:**
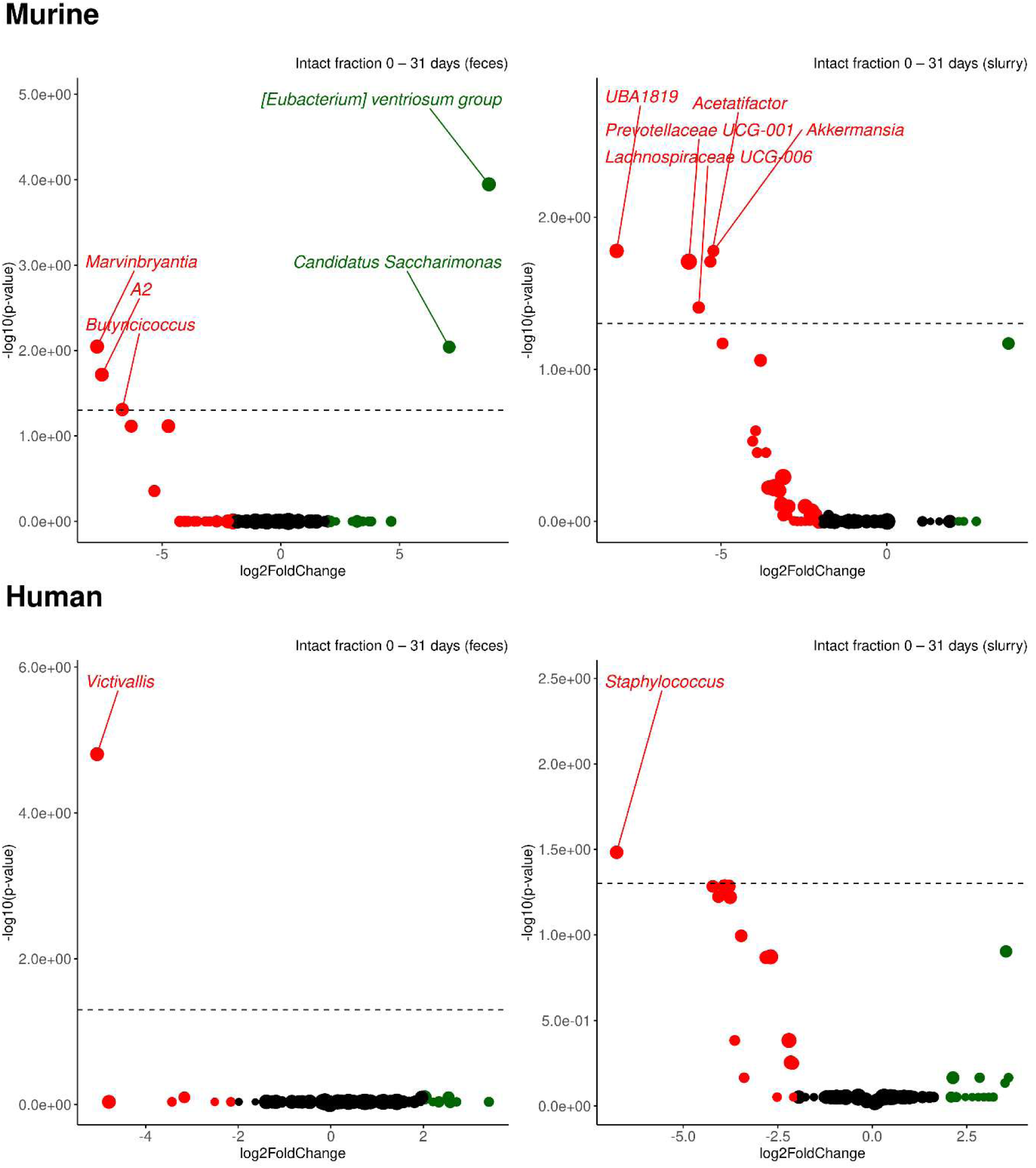
The intact sorted microbiota derived from stored human and murine feces was least affected by direct –80 °C storage of the feces compared to storage as a 5 % glycerol – 1X Tris-EDTA slurry after 31 days. Samples were sequenced with Illumina V3-V4 amplicon 16S rRNA gene sequencing. The intact microbiota was obtained through sorting of SYBR® Green I-positive and Propidium Iodide-negative microbial cells. Storage and sorting were performed in duplicate (except for murine stored feces). Significant ASVs above the dotted line with adjusted p-values <0.05 (Wald tests) were identified with DESeq2. ASVs with a positive shrunken log2 fold change (LFC) were enriched in stored feces (left) or slurry (right) compared to the benchmark (green), whereas ASVs with a negative shrunken LFC were depleted (red). The marker size is proportional to the mean normalized counts.

**Figure 8:**
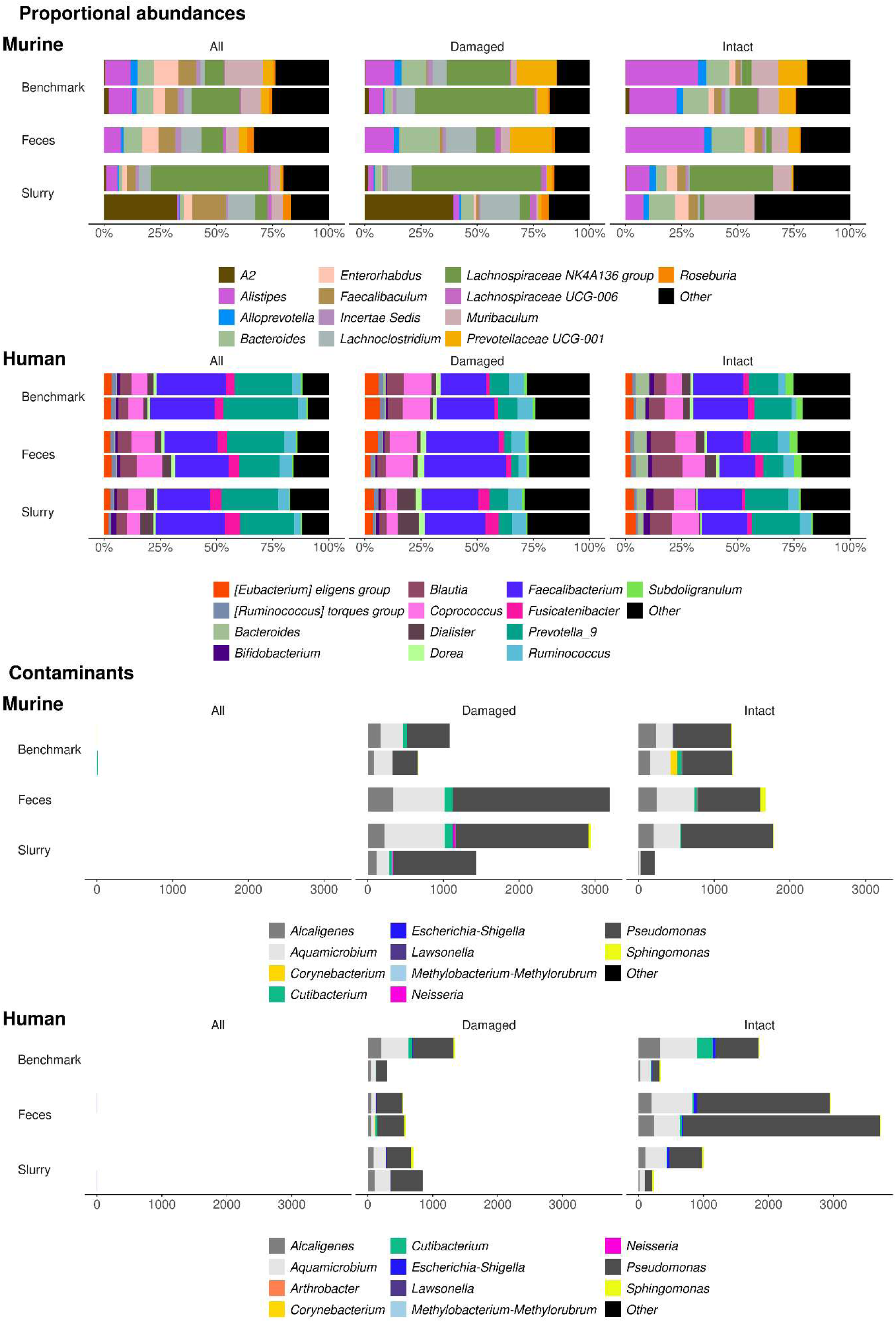
Contaminants were detected in and removed from the metataxonomic sequencing data of sorted microbiota from freshly analyzed (benchmark) and –80°C stored human and murine fecal samples. *In silico* decontaminated sequencing data of directly stored feces was more similar to the benchmark compared to stored fecal slurries containing 5 % glycerol – 1X Tris-EDTA after 31 days. Samples were sequenced with Illumina V3-V4 amplicon 16S rRNA gene sequencing. The intact and damaged microbiota was obtained through sorting of SYBR® Green I– and Propidium Iodide-stained microbial cells. Storage and sorting were performed in duplicate (except for murine stored feces).

Less variability was observed in two different aliquots of human feces. The impact of storage on the intact microbial community was small with a significant decrease in *Victivallis* in directly stored feces (LFC = 1.1, p = 1.6 * 10^-5^) and in *Staphylococcus* in the stored glycerol-TE slurry (LFC = –6.8, p = 3.3 * 10^-2^; Figure 7). The small impact was also reflected in higher community similarities of 61.6 ± 9.1 % between benchmark and stored fecal slurry and 69.2 ± 7.1 % between benchmark and stored feces (Figure S8). The increased similarity resulted from a better preserved viability. The intact cell ratio in human feces dropped from 51.1 % ± 0.1 % to 47.4 % ± 0.1 % in stored feces and to 44.2 % ± 0.0 % in stored fecal slurry.

The FACSome contaminants *Alcaligenes*, *Aquamicrobium* and *Pseudomonas* were detected in the sorted human and murine microbiota (Figure 8). The obtained V3-V4 sequences mapped to the full-length 16S rRNA gene sequences inferred in our two previous experiments (ASV118, 73 and 39, respectively). The previously identified human-skin derived *Cutibacterium* contamination was observed along with *Corynebacterium* which is also part of the skin microbiome. Human manipulation also introduced *Neisseria,* a common salivary microorganism that was formerly identified as a contaminant, in some of the murine sorted samples. The human and murine sorted microbiota were also contaminated by ASVs that mapped to the lab strains *Arthrobacter gandavensis* (94.75 % identity and a BLAST® E-value of 0) and *Escherichia coli str. K-12* substrain MG1655 (100 % identity and a BLAST® E-value of 0).

The recurring detection of FACSome, human-derived and lab contaminants, in three different experiments, performed by three different researchers, at different moments in time, reinforces the need for adequate controls and *in silico* decontamination.

## Discussion

Fluorescence-activated cell sorting (FACS) is a powerful non-destructive, high-throughput, single-cell resolution technique which is often used together with 16S rRNA gene sequencing to probe functionality and activity in microbial subpopulations. The sorted low biomass subpopulations are prone to contamination and bias introduced during sample storage, FACS and DNA extraction. We systematically characterized the widely ignored FACS-sequencing biases for the first time and our insights led to an optimized FACS-sequencing workflow which minimized potential sources of interferences.

The main interference was contamination of the sorted microbiota by *Pseudomonas veronii*, *Aquamicrobium* sp. and *Alcaligenes faecalis.* The hereinafter termed FACSome contaminants were consistently and uniquely found in sorted samples and FACS sheath fluid. FACS sheath fluid surrounds the sample core and focuses the sample into a single-file stream for laser interrogation. After laser interrogation, the sample stream and sheath fluid are mixed trough vibration, enabling dissemination of contaminants in uniform sorting droplets. Contaminated droplets containing target cells are deflected into the sorted cell stream, whereas cell-free droplets are collected as sorted noise. Detection of FACSome species in both the cell and noise fraction suggests that at least part of the contamination stemmed from relic DNA. Relic DNA is stable (Torti et al., 2015) and can persist in the sorter by sticking to the tubing, resulting in carry-over contamination. Relic DNA can also transfer from the sheath fluid tank by passing through the 0.2 µm fluid filter on the sheath fluid line that is connected to the sorting unit (Müller & Nebe-von-Caron, 2010). Relic DNA contamination in the sheath fluid tank can originate from commercially purchased FACS sheath fluid. Alternatively, DNA can be released through cell death, possible induced by sodium fluoride in FACS sheath fluid, of invading environmental microorganisms that are able to grow and possibly form biofilms in the sheath fluid tank (Box et al., 2022; Muraki et al., 2012). All FACSome species are known for their high metabolic diversity and capability to form biofilms (Bhatt et al., 2023; Mustaq et al., 2024; Shoda, 2020). Biofilms can also grow in the sample line and result in contamination through the detachment of cells. While the fluidics system was cleaned daily with ethanol and weekly with bleach solution, *Alcaligenes* and *Pseudomonas* biofilms can resist common disinfectants, especially given the short contact time during run-through cleaning (Chapman, 1998; Mann & Wozniak, 2012; Masák et al., 2014; Peix et al., 2009; Silby et al., 2011; Suwa et al., 2013).

Standard cleaning practices for flow cytometry, including regular bleach disinfection and sheath fluid filtering, could not prevent the introduction of FACSome contaminants (Müller & Nebe-von-Caron, 2010). Future experiments could improve on standard cleaning practices by autoclaving the sheath fluid tank. Autoclaving, however, does not entirely degrade DNA (Calderón-Franco et al., 2020; Gefrides et al., 2010) and cannot be performed on the tubing in the fluidics system. To prevent relic DNA contamination, FACS and DNA extraction can be performed in a HEPA-filtered clean room using bleached and UV-radiated instruments, surfaces and reagents (Stepanauskas et al., 2017). While this is standard practice in single-cell genomics, this is laborious and unavailable in most flow cytometry laboratories. Downstream molecular analysis of the sorted samples, moreover, requires additional cleanrooms, personal protective equipment and robotic handlers to prevent the introduction of carry-over and laboratory contaminants.

Carry-over from high biomass samples was the second most important source of contamination observed in the diluted low biomass samples but could be prevented in the sorted low biomass samples by handling them separated in space and time from the high biomass samples.

Thirdly, sorted low biomass samples were prone to downstream contamination by human skin microorganisms such as *Cutibacterium* and *Corynebacterium*, coming from human sample manipulation (Salter et al., 2014). Sample manipulation also introduced laboratory contaminants, which are effectively disseminated by dust particles and aerosols, especially in BSL-2 facilities where ventilation is limited due to safety restrictions (Weyrich et al., 2019). Other contaminants, such as *Pseudomonas* sp., are commonly found in commercial molecular biology kits, and are referred to as the kitome (Olomu et al., 2020).

Since complete sterilization and DNA decontamination of the laboratory environment is unattainable in standard research facilities, controls should be included to track and remove contaminants in FACS-sequencing data. We used a mock community standard and high biomass reference samples together with anaerobic PBS (aPBS) and FACS sheath fluid blanks to distinguish the FACSome from carry-over, human-derived, kitome, and other laboratory contaminants.

We developed an *in silico* data decontamination procedure to remove all contaminants that did not originate from carry-over and thus did not exceed a 0.1 % abundance in high biomass samples. *In silico* decontamination removed the FACSome and other experiment-specific contaminants, resulting in an increased similarity of sorted and diluted samples to a mock standard. The approach was validated in diverse microbial ecosystems in three separate experiments, performed by three different researchers, at three different moments in time, with two different sequencing techniques. While the FACSome consistently comprised *Alcaligenes faecalis*, *Aquamicrobium* sp. and *Pseudomonas veronii*, variation was observed in the form of an additional unidentified *Bacillus* sp. detected in one of the experiments. Similarly, some variation occurred in the skin-derived and laboratory contaminants and short-read Illumina 16S rRNA gene sequencing indicated a higher abundance of *Cutibacterium* contamination compared to full-length PacBio 16S rRNA gene sequencing. This increased *Cutibacterium* abundance stems from the enhanced amplification of highly fragmented *Cutibacterium* remains from the human skin by Illumina sequencing of the 10-fold smaller V3-V4 region compared to full-length 16S rRNA gene PacBio sequencing. Contaminants are thus experiment-specific and can be safely assumed to be laboratory-specific. As a consequence, the *in silico* decontamination of FACS-sequencing data is experiment– and laboratory-specific. The filters and thresholds applied in the current study should not be used to decontaminate FACS-sequencing data obtained in other studies as subjective prevalence– or abundance-based thresholds set to filter out rare taxa (McMurdie & Holmes, 2013) can remove keystone or indicator taxa (Schloss, 2023; Taguer et al., 2021). Objective prevalence thresholds based on blanks are implemented in the *decontam* R package and the GRIMER decontamination tool (Davis et al., 2018; Piro & Renard, 2023). These tools assume an inverse relationship between the abundance of a contaminant and the sample DNA concentration. A stronger relationship indicates a larger probability that a sequence is a contaminant resulting in a smaller p-value. Based on the default p < 0.1 setting, *Pseudomonas veronii* and *Alcaligenes* sp. were flagged as potential contaminants in our study. FACSome contaminant *Aquamicrobium* sp. remained unchallenged. This indicates that these tools are not suitable to decontaminate FACS-sequencing data. FACS introduced additional contaminants which are absent in high biomass references and thereby differs from dilution, which simply decreased the signal to noise ratio by amplifying the noise which was present but remained undetected in high biomass samples. Sorting, moreover, removes relic DNA and non-targeted microorganisms resulting in true variation compared to high biomass references, which is not accounted for in the underlying *decontam* statistical model. We, therefore, propose an i*n silico* FACS-sequencing decontamination based on the careful inspection of aPBS and FACS sheath fluid blanks to flag potential contaminants and a high biomass reference to exclude carry-over taxa from the flagged contaminants. This workflow based on experiment– and laboratory-specific controls improves the accuracy of microbial community analyses and increases the power of statistical hypothesis testing by reducing the multiplicity problem through the removal of contaminant taxa.

To refine the comparison of sorted subpopulations with a functionality of interest with the bulk microbial community, it is advisable to include samples that are diluted to a low biomass before DNA extraction to account for sequencing bias in low biomass samples.

The comparison of low and high biomass microbial communities was biased by increased input DNA concentrations and PCR amplification used during sequencing library preparation. An increased amplicon product yield resulted in a larger sequencing read depth and an increased observed species richness in various ecosystems. This increased detection of rare taxa with increased DNA input reduced the similarity between high and low biomass microbial communities. The reduced similarity was not driven by subsampling bias upon DNA extraction of low biomass samples, since diluted biomass samples that were amplified with three additional cycles and loaded onto an Illumina flow cell with a six-fold increased input volume, displayed a higher richness. The subsampling effects during sequencing were, moreover, larger than the subsampling effects upon extraction of 10^5^ instead of 10^8^ cells mL^-1^ since read counts are usually below 10^5^. Consequently, our optimized PowerSoil® Pro kit protocol, using a higher sample input volume and bead beating frequency, did not bias microbial community analysis in low biomass compared to high biomass samples.

The optimized PowerSoil® Pro kit protocol with increased bead beating did not demonstrate species-specific DNA extraction bias when applied to a mock community standard and showed an adequate representation of tough-to-lyse G+ species of the *Bacillota* phylum in the fecal samples. A stronger bead beating increases cell lysis, especially for G+ bacteria, and releases more DNA, but could result in increased fragmentation and heat, thereby negatively affecting the DNA quality (Atwood, 2016; Fujimoto et al., 2004; Lyalina et al., 2021; Stepanauskas et al., 2017; Yu, 2016). Fragmentation of DNA due to intensified bead beating was not evident when comparing short-read Illumina or full-length PacBio 16S rRNA gene sequencing data. The data corresponded well, aside from the enrichment of *Cutibacterium* contamination in the Illumina data. This enrichment was likely caused by fragmentation of *Cutibacterium* extracellular relic DNA due to exposure to the inhospitable skin environment.

Relic DNA is persistent in the environment and detected by sequencing (Lennon et al., 2018). While sorting creates a single-cell suspension enabling the selection of sorting droplets with targeted cells of interest, relic DNA can contaminate sorted droplets and interfere with activity-, viability-, or function-targeted single-cell labelling and sorting. This interference was minimal as spiked *Veillonella atypica* relic DNA was effectively removed from the collected mock cell stream (> 99 %) by sorting.

Treatment of samples with AMPure XP beads prior to sorting completely eliminated *V. atypica* relic DNA but the AMPure XP clean-up was not selective to relic DNA and caused major shifts in the microbial community composition. These shifts indicate interactions of cells with the carboxyl surface of the beads that are conventionally used to purify DNA instead of mixtures of cells and DNA (Chen et al., 2020).

A Benzonase digest selectively but not entirely removed spike-in and mock relic DNA. Total relic DNA removal can be pursued by further optimizing the Benzonase incubation which did not involve mixing. Benzonase incubation with mixing has been shown to eliminate relic and host DNA from skin microbiota samples without the introduction of other microbial contamination. This confirmed that the increased contamination with FACSome reads observed in FACS-sequencing data did not stem from Benzonase itself but was due to the combined effect of a decreased amplification of 16S target sequences due to relic DNA degradation and an increased amplification of non-target contaminant DNA introduced during FACS. The decreased target to non-target or signal to noise ratio has been suggested to lie at the basis of the increased richness and detection of rare contaminant taxa in diluted low biomass skin, saliva and upper respiratory tract microbiota (Amar et al., 2021; Biesbroek et al., 2012). We observed minor contamination in the diluted samples and richness, instead, depended on the sequencing library preparation. Differences in library preparation together with relic DNA filtering resulted in dissimilarity between high biomass and non-selectively sorted samples. This dissimilarity is unbiased and relic DNA removal during sorting, possibly enhanced by a Benzonase digest, is vital to ensure accurate activity-, viability-, or function-targeted FACS-sequencing omics analysis.

Activity– and viability-based FACS-sequencing additionally require the preservation of cell integrity during sample storage, which is common in clinical settings. Direct freezing of feces at –80 °C better preserved viability and community composition of FACS-sequenced intact sorted microbiota compared to glycerol-TE cryoprotectant amended fecal slurries. Strictly anaerobic *Lachnospiraceae* were most sensitive to storage of murine fecal samples which were not processed anaerobically. Anaerobic processing of fecal samples before freezing might increase the viability of strict anaerobes, but is laborious and unpractical in clinical settings (Papanicolas et al., 2019). Alternative cryoprotectants such as sucrose and inulin have been shown to improve the preservation of viable cultured strict anaerobes (Bircher et al., 2018). However, the effects of storage and freezing are species– and sample matrix-specific. It is, therefore, advisable to optimize storage conditions specifically for the investigated microbial ecosystem.

## Conclusion

Bulk meta-omics fail to provide sufficient functional resolution to gain mechanistic insights in microbial ecosystems. Function-driven microbiome research through fluorescence-activated cell sorting (FACS) combined with function-, activity– or viability-targeted labeling results in low biomass samples, presenting a currently overlooked challenge for accurate metataxonomic profiling. We identified sheath fluid, human skin-derived and laboratory contaminants introduced during FACS-sequencing and we developed a protocol to *in silico* decontaminate the sequencing data based on co-analyzed sheath fluid and PBS blanks and a high biomass reference. We advise to separate high and low biomass sample manipulation in space and time and to use the PowerSoil Pro kit with increased bead beating frequency and sample input volume for DNA extraction. A Benzonase treatment is optional to optimize relic DNA removal and we recommend direct –80°C sample storage of murine and human feces to minimize bias in the viable sorted microbiota. Our optimized extraction and decontamination workflow improves the accuracy of metataxonomic FACS-sequencing analysis and is hypothesized to advance other sequencing-based omics analyses of sorted microbial subpopulations. Our optimized FACS-sequencing workflow is thus imperative to unlock the full potential of function-driven single-cell labelling and FACS, in turn enabling mode-of-action studies that will broaden our understanding of microbe-environment interactions.

## Acknowledgements

The research was funded by the Special Research Fund (BOF) Concerted Research Actions (GOA, grant number: 01G03122) from the Flemish government and by the Research Foundation – Flanders (FWO, Belgium) (grant number: G0B2719N). N.V. is funded by a PhD fellowship strategic basic research (FWO, grant number: 1S09825N). The authors would like to thank Tim Lacoere for the technical support and Hannah Lernout of the Department of Internal Medicine and Pediatrics (Ghent University, Ghent, Belgium) for the murine samples.

---

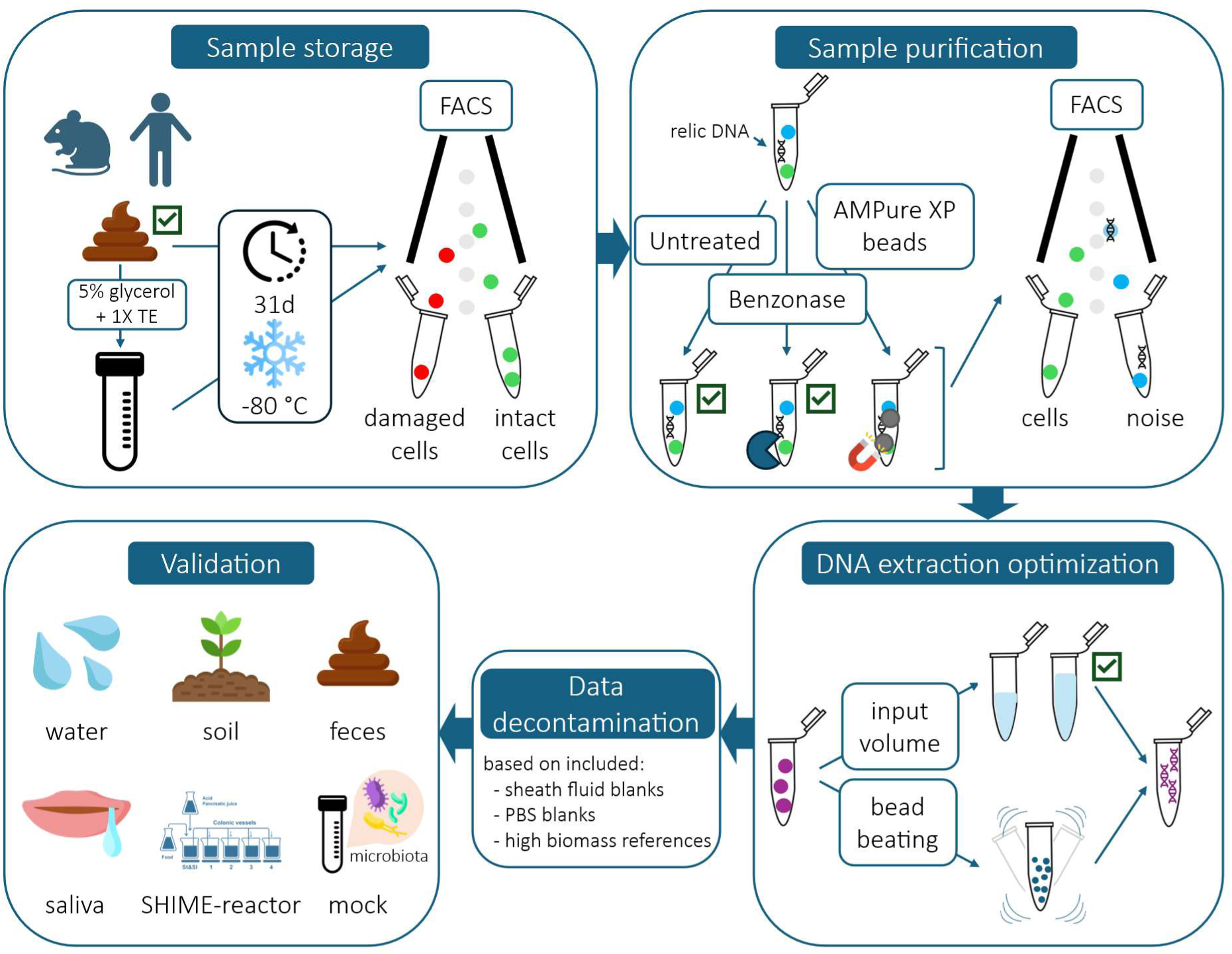

