## Supplementary material for "An optimized flow cytometry sorting-sequencing workflow reduces storage, sorting and extraction bias in sorted microbial communities"

### Supplementary figures

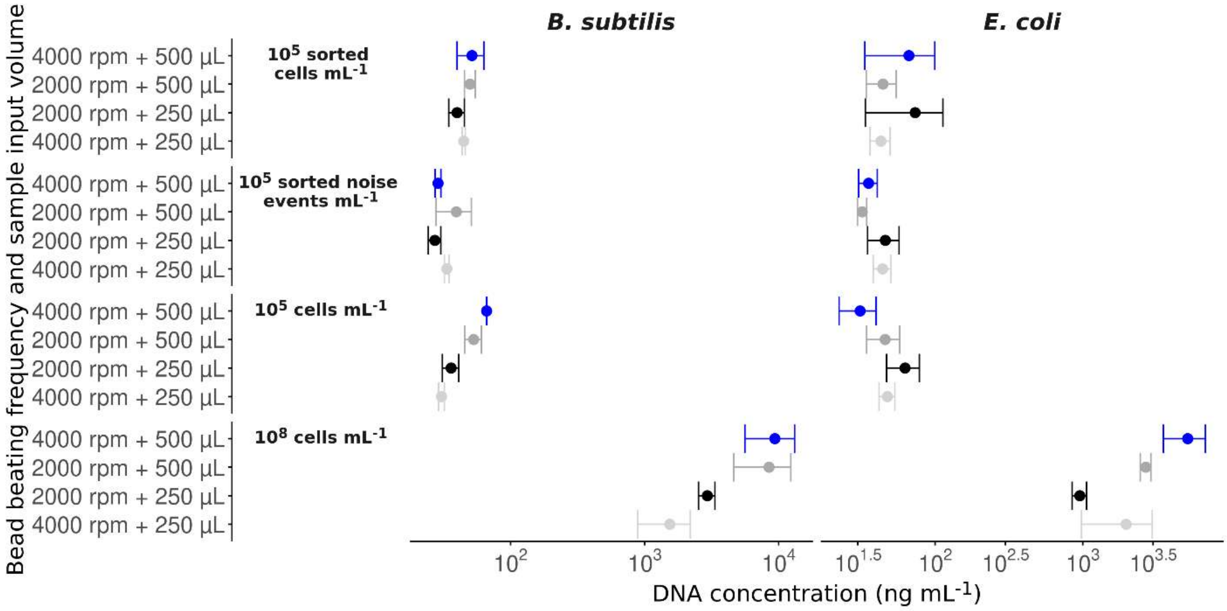


**Figure S1**: The PowerSoil® Pro protocol with an increased 500 µL input volume and 4,000 rpm bead beating (blue) yielded the highest average (n=2) double-stranded DNA quantity (ng mL$^{-1}$) determined by the QuantiFluor® dsDNA System in sorted (~ 2.88 * 10$^{5}$ cells mL$^{-1}$), diluted (~ 10$^{5}$ cells mL$^{-1}$) and high biomass reference (~ 10$^{8}$ cells mL$^{-1}$) samples and sorted noise (~ 2.88 * 10$^{5}$ noise events mL$^{-1}$) derived from pure cultures of *Bacillus subtilis* subsp. *subtilis* strain 168 and of Escherichia coli str. K-12 substr. MG1655. The other protocols using 500 µL at 2,000 rpm (dark gray) and 250 µL at 4,000 rpm (light grey) and the reference (black) performed equally good or better to extract sorted noise (~ 2.88 * 10$^{5}$ noise events mL$^{-1}$).

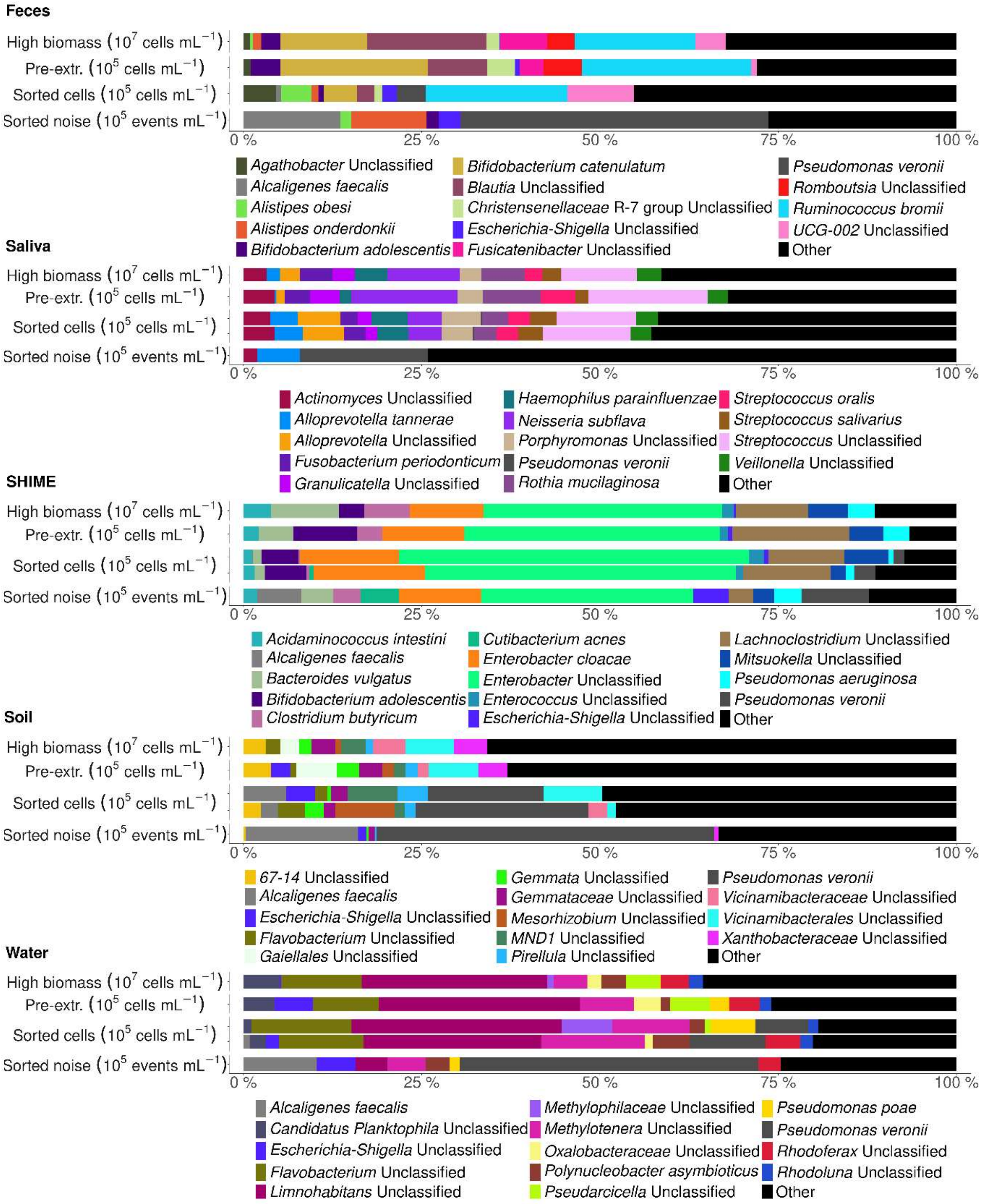


**Figure S2:** Contaminants were detected in the metataxonomic sequencing data of low biomass feces, saliva, SHIME gut reactor, soil and water microbiota. The removal of relic DNA during sorting resulted in discrepancies between the non-sorted and sorted samples. Samples were sequenced with PacBio full-length 16S rRNA gene sequencing. The low biomass was obtained through dilution of the high biomass reference samples ($10^7$ cells mL$^{-1}$) to $10^5$ cells mL$^{-1}$ before extraction (Pre-extr.) and through sorting of the high biomass samples into $10^5$ cells mL$^{-1}$ and $10^5$ noise events mL$^{-1}$ using a non-selective SYBR® Green I staining. Dilution and sorting was performed in duplicate (except for feces).

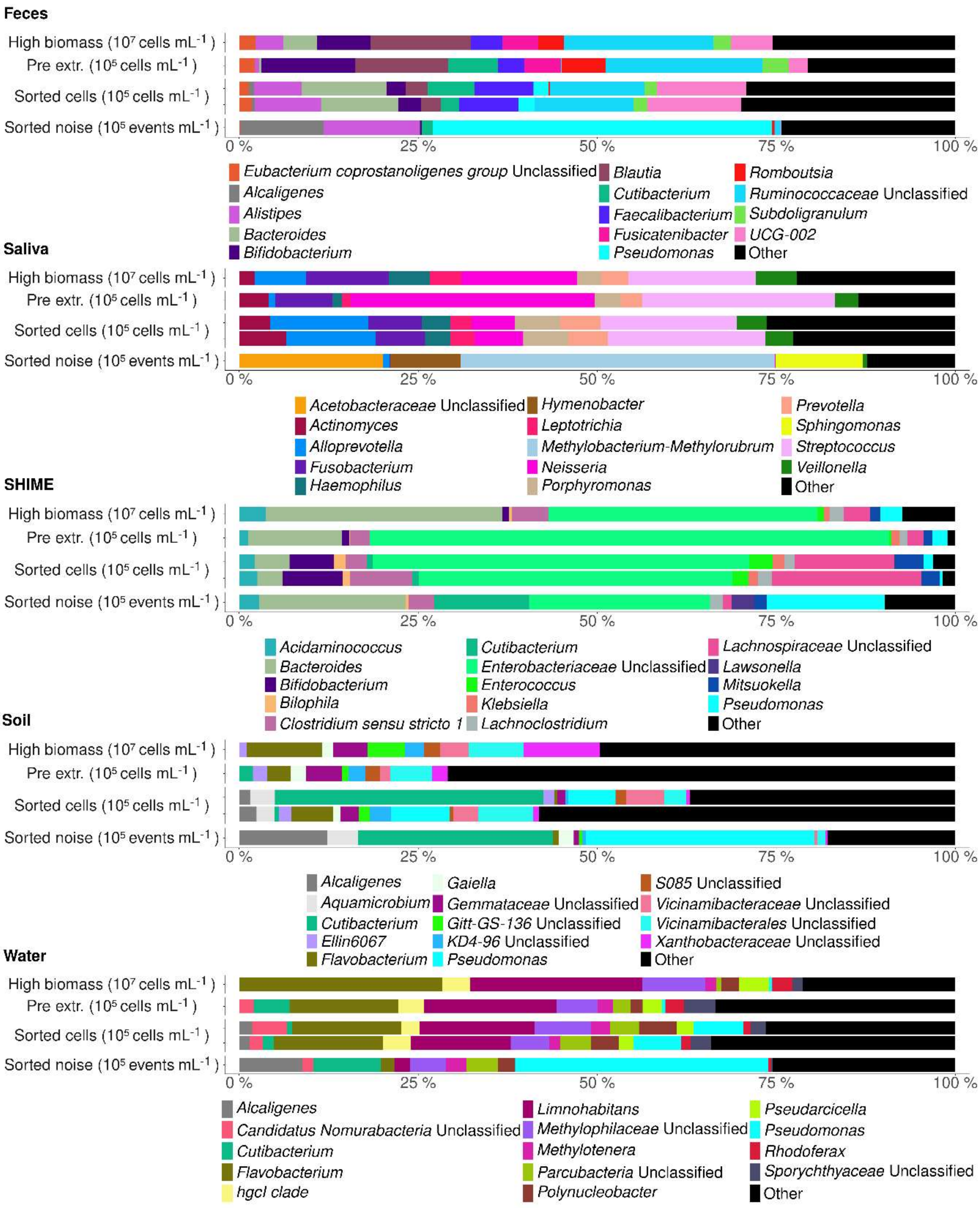


**Figure S3:** Contaminants were detected in the metataxonomic sequencing data of low biomass feces, saliva, SHIME gut reactor, soil and water microbiota. The removal of relic DNA during sorting resulted in discrepancies between the non-sorted and sorted samples. Samples were sequenced with Illumina V3-V4 amplicon 16S rRNA gene sequencing. The low biomass was obtained through dilution of the high biomass reference samples ($10^7$ cells $mL^{-1}$) to $10^5$ cells $mL^{-1}$ before extraction (Pre-extr.) and through sorting of the high biomass samples into $10^5$ cells $mL^{-1}$ and $10^5$ noise events $mL^{-1}$ using a non-selective SYBR® Green I staining. Dilution and sorting was performed in duplicate (except for feces).

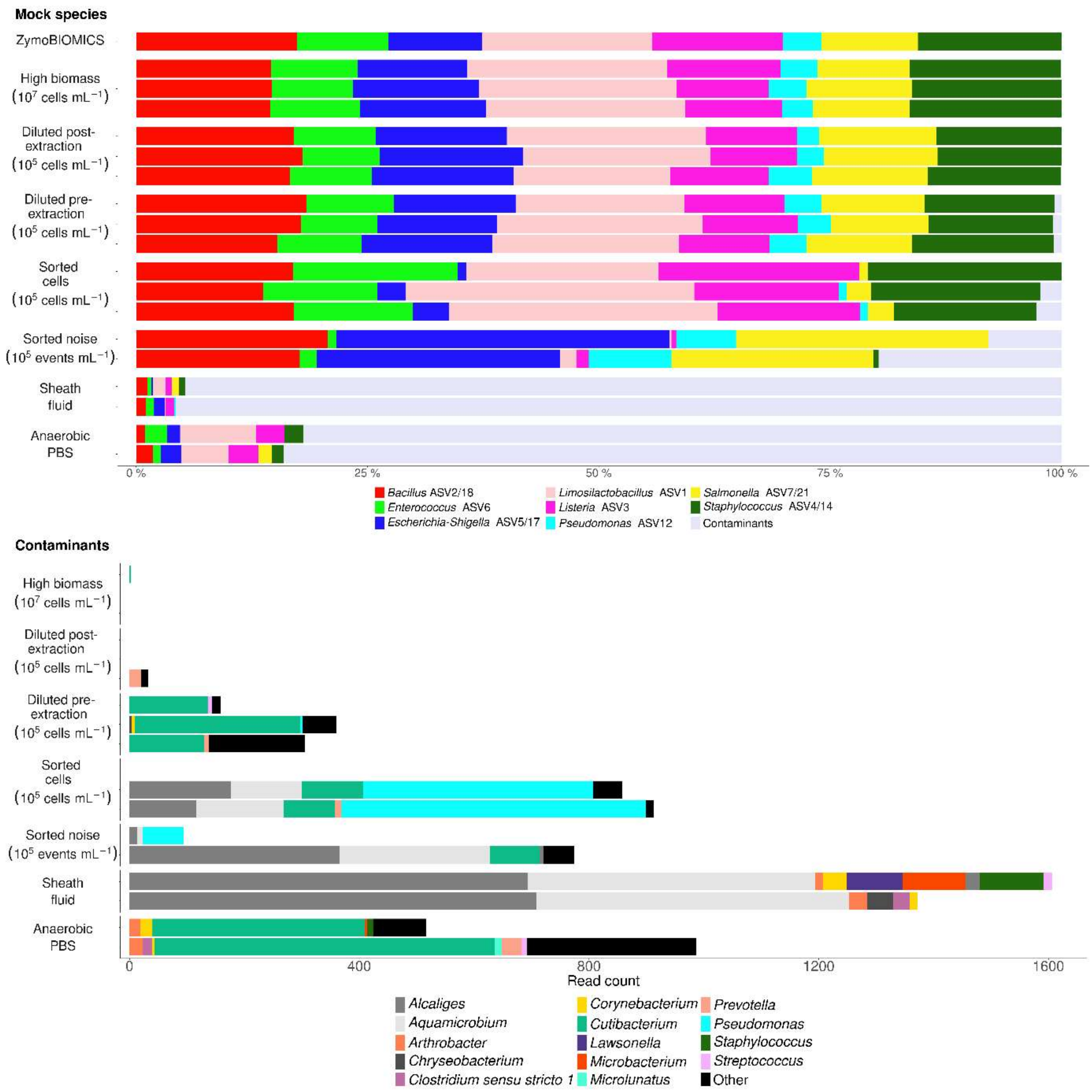


**Figure S4:** Contaminants were detected in the metataxonomic sequencing data of low biomass ZymoBIOMICS mock microbiota. Samples were sequenced with Illumina V3-V4 amplicon 16S rRNA gene sequencing. The low biomass was obtained through dilution of the high biomass mock standard ($10^7$ cells $mL^{-1}$) to $10^5$ cells $mL^{-1}$ before (diluted pre-extraction) and after (diluted post-extraction) DNA extraction and through sorting of the mock standard into $10^5$ cells $mL^{-1}$ and $10^5$ noise events $mL^{-1}$ using a non-selective SYBR® Green I staining. Dilution and sorting were performed in triplicate. FACS sheath fluid and aPBS diluent blanks were included in duplicate to identify and track the FACSome contaminants introduced during FACS-sequencing.

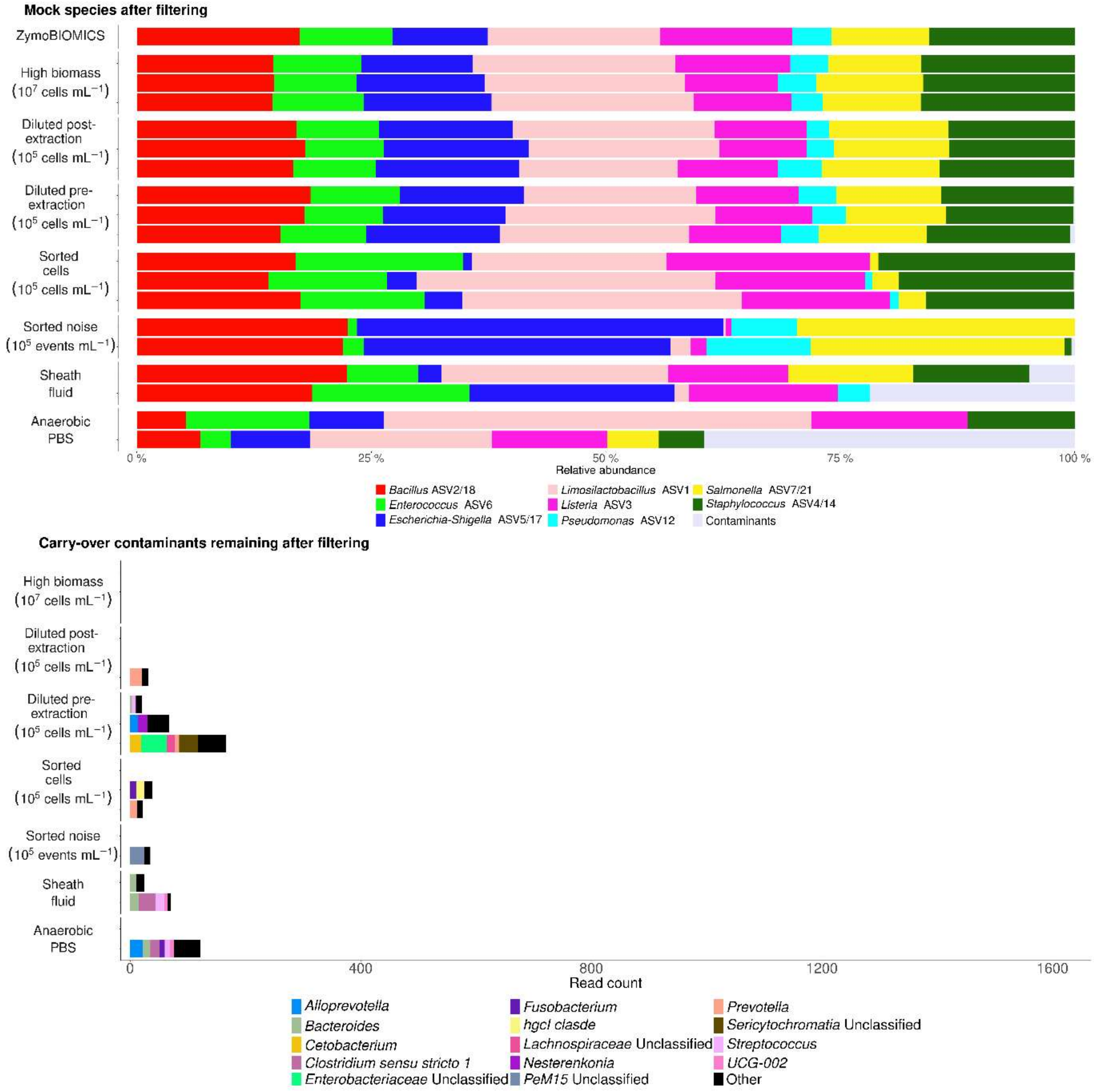


**Figure S5:** *In silico* data decontamination removed contaminants from the metataxonomic sequencing data of low biomass ZymoBIOMICS mock microbiota. The removal of relic DNA during sorting resulted in discrepancies between the non-sorted and sorted samples. Samples were sequenced with Illumina V3-V4 amplicon 16S rRNA gene sequencing. The low biomass was obtained through dilution of the high biomass mock standard ($10^7$ cells $mL^{-1}$) to $10^5$ cells $mL^{-1}$ before (diluted pre-extraction) and after (diluted post-extraction) DNA extraction and through sorting of the mock standard into $10^5$ cells $mL^{-1}$ and $10^5$ noise events $mL^{-1}$ using a non-selective SYBR® Green I staining. Dilution and sorting were performed in triplicate. FACS sheath fluid and aPBS diluent blanks were included in duplicate to identify and track the FACSome contaminants introduced during FACS-sequencing.

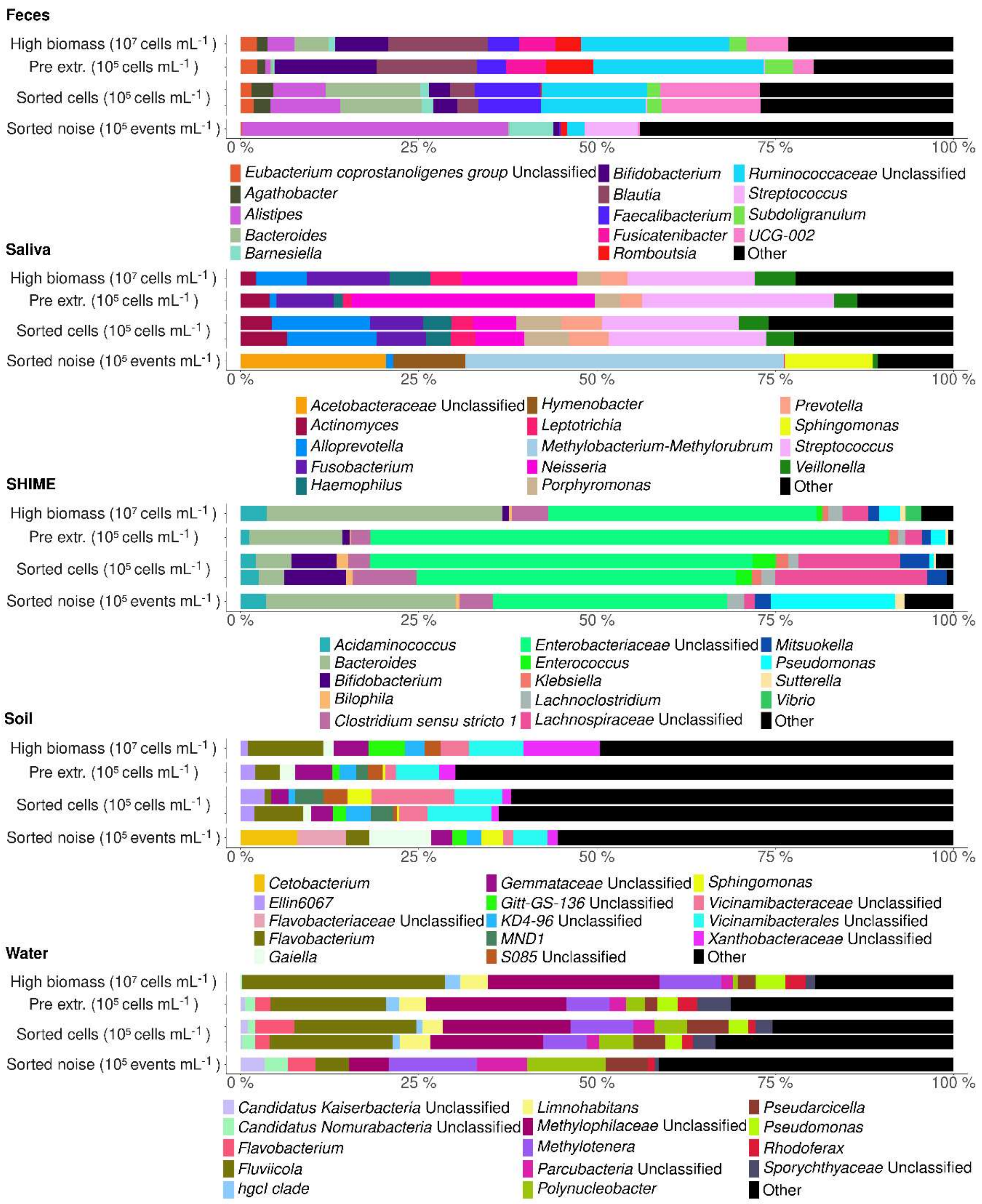


**Figure S6**: *In silico* data decontamination removed contaminants from the metataxonomic sequencing data of low biomass feces, saliva, SHIME gut reactor, soil and water microbiota. The removal of relic DNA during sorting resulted in discrepancies between the non-sorted and sorted samples. Samples were sequenced with Illumina V3-V4 amplicon 16S rRNA gene sequencing. The low biomass was obtained through dilution of the high biomass reference samples ($10^7$ cells mL$^{-1}$) to $10^5$ cells mL$^{-1}$ before extraction (Pre-extr.) and through sorting of the high biomass samples into $10^5$ cells mL$^{-1}$ and $10^5$ noise events mL$^{-1}$ using a non-selective SYBR® Green I staining. Dilution and sorting were performed in duplicate (except for feces).

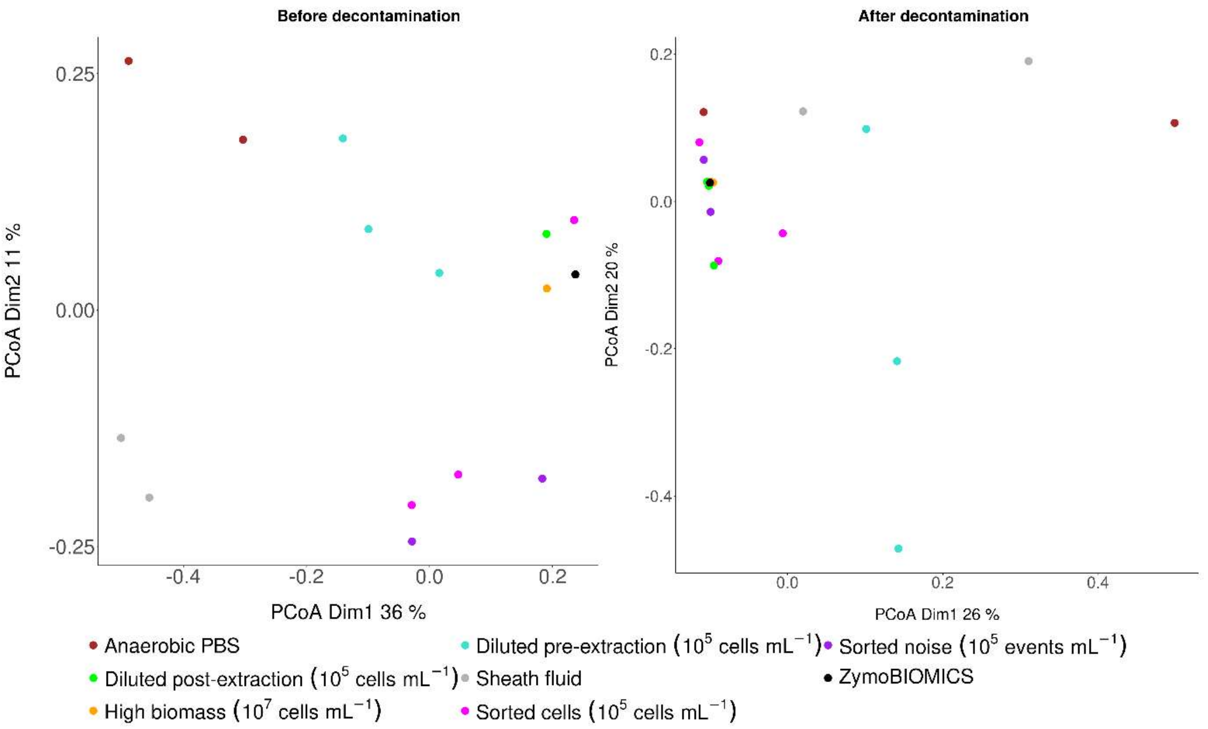


**Figure S7**: *In silico* data decontamination increased the absence/presence-based Jaccard similarity between sorted ZymoBIOMICS mock cells ($10^5$ cells $mL^{-1}$; pink) and sorted noise events ($10^5$ $mL^{-1}$; purple) and the theoretical ZymoBIOMICS standard (black) resulting in a closer ordination of these samples in a Principal Coordinates Analysis (PCoA). Low biomass samples obtained through dilution of the high biomass mock standard (orange; $10^7$ cells $mL^{-1}$) to $10^5$ cells $mL^{-1}$ before (diluted pre-extraction; turquoise) and after (diluted post-extraction; green) DNA extraction did not display an improved clustering. Samples were sequenced with Illumina V3-V4 amplicon 16S rRNA gene sequencing. Dilution and sorting with a non-selective SYBR® Green I cell staining were performed in triplicate. FACS sheath fluid (light grey) and aPBS (brown) diluent blanks were included in duplicate to identify and track the FACSome contaminants introduced during FACS-sequencing.

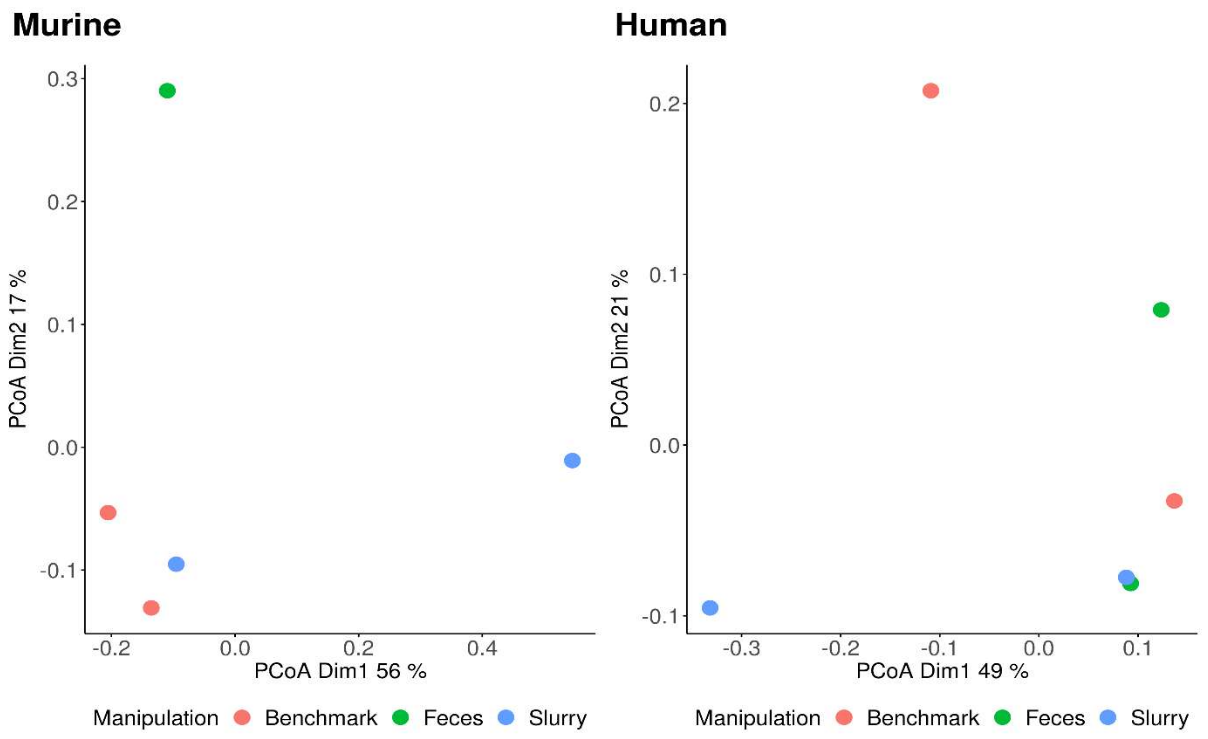


**Figure S8**: The absence/presence-based Jaccard similarity of the intact microbiota, calculated based on *in silico* decontaminated Illumina V3-V4 amplicon 16S rRNA gene sequencing data, was less similar between stored murine fecal samples and freshly analyzed benchmarks compared to the stored human fecal samples and freshly analyzed benchmarks after 31 days. Slurries contained 5 % glycerol - 1X Tris-EDTA. SYBR® Green I- and Propidium Iodide-stained fecal samples were sorted into intact and damaged fractions based on a heat-killed control. Samples were stored in duplicate.

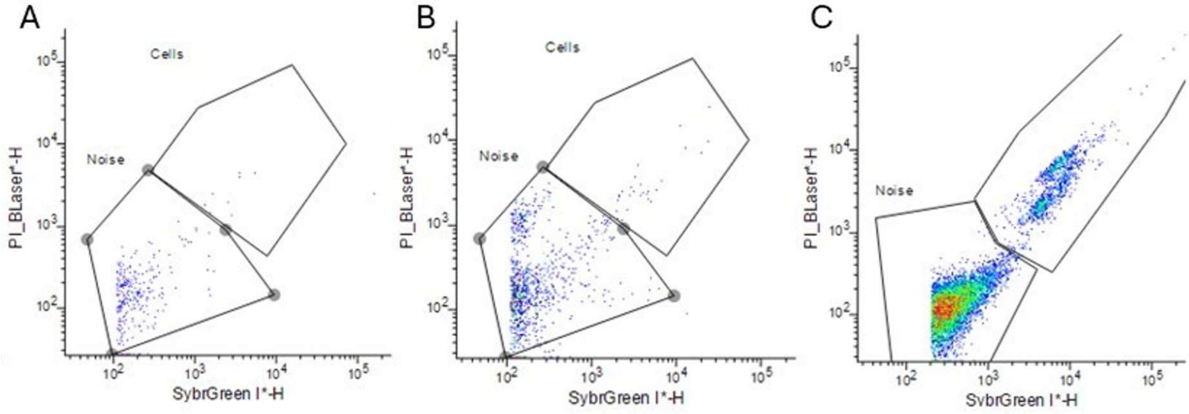


**Figure S9:** Samples stained with SYBR® Green I were sorted into a SYBR green-positive cell fraction and a cell-free noise fraction based on the gating of blue laser biplots in the BD FACSChorus$^{TM}$ software. Gating was based on a 0.2 µm filtered sample (A) and aPBS (B) for cell-free noise and a sample of the mock community (C) for SYBR® Green I-positive cells.

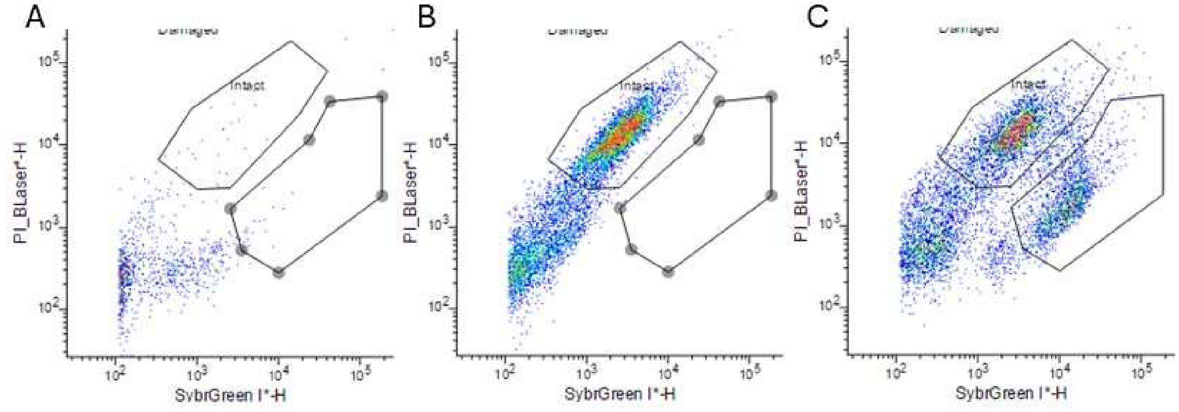


**Figure S10:** Samples stained with SYBR® Green I and Propidium Iodide were sorted into an intact cell fraction and a damaged cell fraction based on the gating of blue laser biplots in the BD FACSChorus$^{TM}$ software. Gating was based on a 0.2 µm filtered sample (A) for cell-free noise, a heat-killed sample (B) for the SYBR® Green I-positive and Propidium Iodide-positive damaged cells and a fecal sample (C) for SYBR® Green I-positive intact cells.

### Supplementary tables

**Table S1**: An increased DNA input (ng) and PCR amplification (33 cycles) were used during sequencing library preparation of low biomass samples for Illumina V3-V4 amplicon 16S rRNA gene sequencing compared to PacBio full-length 16S rRNA gene sequencing (20 cycles). High biomass samples were PCR amplified with 30 cycles for Illumina 16S rRNA gene sequencing and with 20 cycles for PacBio full-length 16S rRNA gene sequencing. An increased DNA input and PCR amplification resulted in increased amplicon product yields, in turn resulting in a larger sequencing read depth (read counts). The low biomass was obtained through dilution of high biomass ZymoBIOMICS mock community, feces, saliva, SHIME gut reactor, soil and water samples ($10^7$ cells $mL^{-1}$) to $10^5$ cells $mL^{-1}$ before (diluted pre-extraction) and after (diluted post-extraction) DNA extraction and through sorting of the high biomass samples into $10^5$ cells $mL^{-1}$ and $10^5$ noise events $mL^{-1}$ using a non-selective SYBR® Green I staining. FACS sheath fluid and aPBS diluent blanks were included in duplicate. DNA concentrations in DNA extracts (ng/µL) were measured with a Qubit Flex fluorometer (Invitrogen) using the Qubit™ 1X dsDNA High Sensitivity Assay Kit by VIB Nucleomics Core (Leuven, Belgium).

| Microbial ecosystem | Sample | DNA concentration in DNA extract (ng/µL) | DNA input for sequencing library preparation (ng) | | Read counts | |
|---|---|---|---|---|---|---|
| | | | PacBio | Illumina | PacBio | Illumina |
| Mock | High biomass | 0.176 | 1.760 | 0.176 | 123459 | 60953 |
| Mock | High biomass | 0.672 | 6.720 | 0.672 | 117471 | 38974 |
| Mock | High biomass | 0.376 | 3.760 | 0.376 | 118693 | 35324 |
| Feces | High biomass | 0.005 | 0.052 | 0.005 | 65042 | 11308 |
| Saliva | High biomass | 0.702 | 7.020 | 0.702 | 122185 | 54681 |
| SHIME | High biomass | 0.922 | 9.220 | 0.922 | 184406 | 66612 |
| Soil | High biomass | 0.922 | 9.220 | 0.922 | 118538 | 20936 |
| Water | High biomass | 0.019 | 0.194 | 0.019 | 68789 | 8941 |
| Mock | Diluted post-extraction | 0.013 | 0.131 | 0.078 | 110611 | 147702 |
| Mock | Diluted post-extraction | 0.036 | 0.356 | 0.216 | 121664 | 173514 |

| | | | | | | |
|---|---|---|---|---|---|---|
| Mock | Diluted post-extraction | 0.022 | 0.224 | 0.132 | 115808 | 100180 |
| Mock | Diluted pre-extraction | 0.003 | 0.022 | 0.018 | 61387 | 124613 |
| Mock | Diluted pre-extraction | 0.002 | 0.011 | 0.012 | 19247 | 163866 |
| Mock | Diluted pre-extraction | 0.002 | 0.016 | 0.012 | 33821 | 75555 |
| Feces | Diluted pre-extraction | 0.016 | 0.111 | 0.096 | 10306 | 24182 |
| Saliva | Diluted pre-extraction | 0.0003 | 0.002 | 0.002 | 78245 | 137570 |
| SHIME | Diluted pre-extraction | 0.018 | 0.089 | 0.108 | 31596 | 16337 |
| Soil | Diluted pre-extraction | 0.029 | 0.202 | 0.174 | 5056 | 11141 |
| Water | Diluted pre-extraction | 0.046 | 0.276 | 0.276 | 4585 | 21742 |
| Mock | Sorted cells | 0.003 | 0.032 | 0.018 | 85203 | 142723 |
| Mock | Sorted cells | 0.013 | 0.107 | 0.078 | 65722 | 134685 |
| Mock | Sorted cells | 0.006 | 0.038 | 0.036 | 59629 | 4057 |
| Feces | Sorted cells | 0.008 | 0.041 | 0.048 | 7956 | 13469 |
| Saliva | Sorted cells | 0.013 | 0.080 | 0.078 | 67939 | 94778 |
| Saliva | Sorted cells | 0.022 | 0.154 | 0.132 | 34024 | 88679 |
| SHIME | Sorted cells | 0.015 | <0.075 | 0.090 | 5279 | 4351 |
| SHIME | Sorted cells | 0.002 | 0.018 | 0.012 | 12430 | 92824 |
| Soil | Sorted cells | 0.026 | 0.132 | 0.156 | 1753 | 56341 |
| Soil | Sorted cells | 0.007 | 0.037 | 0.042 | 1885 | 4602 |
| Water | Sorted cells | 0.006 | 0.030 | 0.036 | 2302 | 18066 |
| Water | Sorted cells | 0.016 | 0.081 | 0.096 | 1910 | 18638 |
| Mock | Sorted noise | 0.007 | 0.034 | 0.042 | 4929 | 43359 |
| Mock | Sorted noise | 0.015 | 0.103 | 0.090 | 1828 | 52 |
| Mock | Sorted noise | 0.001 | 0.005 | 0.006 | 12523 | 4881 |
| Feces | Sorted noise | 0.003 | 0.017 | 0.018 | 1719 | 28650 |
| Saliva | Sorted noise | 0.014 | 0.081 | 0.084 | 2051 | 19998 |
| SHIME | Sorted noise | 0.016 | 0.095 | 0.096 | 2820 | 18934 |

| | | | | | | |
|---|---|---|---|---|---|---|
| Soil | Sorted noise | 0.021 | 0.144 | 0.126 | 4735 | 5460 |
| Water | Sorted noise | 0.002 | <0.012 | 0.012 | 2009 | 9002 |
| Sheath fluid | | 0.007 | 0.069 | 0.042 | 94564 | 27976 |
| Sheath fluid | | 0.006 | <0.031 | 0.036 | 1422 | 41295 |
| aPBS | | 0.017 | 0.149 | 0.102 | 1264 | 7092 |
| aPBS | | 0.017 | 0.171 | 0.102 | 697 | 3114 |

**Table S2:** Low biomass samples sequenced with increased DNA input (ng) and PCR amplification (33 cycles) on an Illumina instrument had an increased observed Chao species richness compared to high biomass samples before and after decontamination, in contrast to PacBio sequenced samples. The low biomass was obtained through dilution of high biomass ZymoBIOMICS mock community, feces, saliva, SHIME gut reactor, soil and water samples ($10^7$ cells $mL^{-1}$) to $10^5$ cells $mL^{-1}$ before DNA extraction.

| **Before decontamination** | **Microbial ecosystem** | **PacBio** | | **Illumina** | |
|---|---|---|---|---|---|
| | | **Low biomass** | **High biomass** | **Low biomass** | **High biomass** |
| | Mock | 30 | 12 | 29 | 12 |
| | Mock | 49 | 10 | 27 | 12 |
| | Mock | 17 | 10 | 18 | 13 |
| | Feces | 66 | 51 | 92 | 92 |
| | Saliva | 46 | 45 | 123 | 98 |
| | SHIME | 15 | 33 | 26 | 58 |
| | Soil | 83 | 471 | 249 | 78 |
| | Water | 58 | 423 | 242 | 57 |
| **After decontamination** | **Microbial ecosystem** | **Low biomass** | **High biomass** | **Low biomass** | **High biomass** |
| | Mock | 29 | 11 | 26 | 12 |
| | Mock | 46 | 10 | 24 | 12 |
| | Mock | 16 | 10 | 16 | 12 |
| | Feces | 60 | 51 | 60 | 51 |
| | Saliva | 45 | 44 | 45 | 44 |
| | SHIME | 15 | 33 | 15 | 33 |
| | Soil | 82 | 471 | 143 | 57 |
| | Water | 56 | 420 | 133 | 39 |

**Table S3**: Contaminant reads and ASVs were detected in sheath fluid blanks (n=2 biological replicates) with PacBio full-length 16S rRNA gene sequencing.

| ASV | Species | Read count of replicates | |
|---|---|---|---|
| ASV16 | *Bifidobacterium adolescentis* | 12 | 0 |
| ASV39 | *Pseudomonas veronii* | 938 | 164 |
| ASV73 | *Aquamicrobium Unclassified* | 0 | 28 |
| ASV101 | *Escherichia-Shigella Unclassified* | 44 | 0 |
| ASV118 | *Alcaligenes faecalis* | 327 | 30 |
| ASV892 | *Ilumatobacter Unclassified* | 2 | 0 |
| ASV1242 | *Cutibacterium acnes* | 7 | 58 |
| ASV1262 | *Pseudomonas poae* | 14 | 0 |
| ASV1272 | *Escherichia-Shigella Unclassified* | 51 | 57 |
| ASV1782 | *Arthrobacter gandavensis* | 187 | 25 |
| ASV2949 | *Curvibacter Unclassified* | 29 | 0 |
| ASV13964 | *Allorhizobium-Neorhizobium-Pararhizobium-Rhizobium Unclassified* | 0 | 11 |
| ASV14029 | *Methylobacterium-Methylorubrum adhaesivum* | 8 | 0 |
| ASV19118 | *Lawsonella Unclassified* | 0 | 7 |

**Table S4**: Contaminant reads and ASVs were detected in aPBS blanks (n=2 biological replicates) with PacBio full-length 16S rRNA gene sequencing.

| ASV | Species | Read count of replicates | |
|---|---|---|---|
| ASV21 | *Bacteroides vulgatus* | 0 | 5 |
| ASV81 | *Bacteroides Unclassified* | 5 | 0 |
| ASV92 | *Megamonas Unclassified* | 14 | 0 |
| ASV101 | *Escherichia-Shigella Unclassified* | 37 | 26 |
| ASV127 | *Enterobacter Unclassified* | 0 | 13 |
| ASV1242 | *Cutibacterium acnes* | 48 | 15 |
| ASV1262 | *Pseudomonas poae* | 60 | 0 |
| ASV1272 | *Escherichia-Shigella Unclassified* | 27 | 50 |
| ASV2949 | *Curvibacter Unclassified* | 25 | 0 |
| ASV6421 | *Pseudomonas poae* | 0 | 49 |
| ASV7715 | *Paracoccus chinensis* | 5 | 0 |
| ASV8686 | *Lawsonella Unclassified* | 5 | 0 |
| ASV10198 | *Haemophilus parainfluenzae* | 16 | 0 |
| ASV13515 | *Candidatus Finniella Unclassified* | 0 | 20 |
| ASV17496 | *Corynebacterium tuberculostearicum* | 10 | 0 |
| ASV18688 | *Sphingomonas melonis* | 8 | 0 |

**Table S5**: Contaminant reads and ASVs were detected in the sorted mock cells and noise (n=3 biological replicates) using PacBio full-length 16S rRNA gene sequencing.

| ASV | Species | Sorted cells (n=3) | | | Sorted noise (n=3) | | |
|---|---|---|---|---|---|---|---|
| | | Reads | Reads | Reads | Reads | Reads | Reads |
| ASV13 | *Bacteroides dorei* | 0 | 0 | 4 | 0 | 0 | 0 |
| ASV17 | *Bifidobacterium adolescentis* | 0 | 4 | 0 | 0 | 0 | 0 |
| ASV39 | *Pseudomonas veronii* | 638 | 213 | 168 | 265 | 72 | 461 |
| ASV50 | *Streptococcus salivarius* | 0 | 0 | 0 | 0 | 0 | 9 |
| ASV73 | *Aquamicrobium Unclassified* | 80 | 26 | 30 | 72 | 0 | 96 |
| ASV91 | *Megamonas Unclassified* | 0 | 0 | 26 | 0 | 0 | 0 |
| ASV101 | *Escherichia-Shigella Unclassified* | 0 | 0 | 0 | 8 | 0 | 0 |
| ASV118 | *Alcaligenes faecalis* | 190 | 67 | 38 | 62 | 0 | 144 |
| ASV181 | *Enterococcus thailandicus* | 3 | 0 | 0 | 0 | 0 | 0 |
| ASV572 | *Prevotella_9 Unclassified* | 6 | 0 | 0 | 0 | 0 | 0 |
| ASV1145 | *Holdemanella Unclassified* | 0 | 7 | 0 | 0 | 0 | 0 |
| ASV1242 | *Cutibacterium acnes* | 0 | 27 | 0 | 24 | 0 | 55 |
| ASV1246 | *Clostridium sensu stricto 1 Unclassified* | 3 | 0 | 0 | 0 | 0 | 0 |
| ASV1262 | *Pseudomonas poae* | 0 | 0 | 27 | 0 | 53 | 12 |
| ASV1681 | *Haemophilus parainfluenzae* | 0 | 0 | 0 | 3 | 0 | 0 |
| ASV1870 | *Rothia dentocariosa* | 0 | 0 | 0 | 10 | 0 | 0 |
| ASV2257 | *Gemella haemolysans* | 6 | 0 | 0 | 0 | 0 | 0 |
| ASV2949 | *Curvibacter Unclassified* | 5 | 0 | 0 | 0 | 22 | 0 |
| ASV3664 | *Pseudomonas poae* | 11 | 0 | 0 | 31 | 0 | 0 |
| ASV4566 | *Streptococcus Unclassified* | 0 | 0 | 0 | 0 | 44 | 0 |
| ASV6410 | *Herbaspirillum huttiense* | 0 | 12 | 0 | 0 | 0 | 0 |

| | | | | | | | |
|---|---|---|---|---|---|---|---|
| ASV7629 | *Herbaspirillum huttiense* | 0 | 0 | 0 | 0 | 6 | 0 |
| ASV7881 | *Neisseria subflava* | 0 | 8 | 0 | 0 | 0 | 0 |
| ASV8307 | *Neisseria subflava* | 7 | 0 | 0 | 21 | 0 | 0 |
| ASV9362 | *Cutibacterium acnes* | 0 | 0 | 0 | 0 | 12 | 0 |
| ASV10081 | *Chryseobacterium anthropi* | 0 | 0 | 0 | 0 | 16 | 0 |
| ASV10811 | *Staphylococcus epidermidis* | 0 | 0 | 0 | 6 | 0 | 0 |
| ASV12451 | *Schlesneria Unclassified* | 0 | 0 | 0 | 24 | 0 | 0 |
| ASV13143 | *Chryseobacterium anthropi* | 0 | 0 | 0 | 0 | 21 | 0 |
| ASV13425 | *Staphylococcus hominis* | 0 | 0 | 0 | 20 | 0 | 0 |
| ASV14646 | *Porphyromonas Unclassified* | 0 | 0 | 16 | 0 | 0 | 0 |
| ASV16195 | *Corynebacterium propinquum* | 0 | 0 | 0 | 0 | 2 | 0 |
| ASV16268 | *Enterobacter cloacae* | 0 | 12 | 0 | 0 | 0 | 0 |
| ASV17277 | *Halomonas Unclassified* | 0 | 0 | 0 | 0 | 10 | 0 |
| ASV19081 | *Pseudomonas mendocina* | 0 | 0 | 0 | 7 | 0 | 0 |
| ASV19801 | *Cutibacterium acnes* | 0 | 0 | 6 | 0 | 0 | 0 |

**Table 6**: Contaminant reads and ASVs were detected in sheath fluid blanks (n=2 biological replicates) using Illumina V3-V4 amplicon 16S rRNA gene sequencing.

| ASV | Genus | Read count of replicates | |
|---|---|---|---|
| ASV10 | *Streptococcus* | 15 | 0 |
| ASV11 | *Pseudomonas* | 2883 | 2410 |
| ASV13 | *Cutibacterium* | 777 | 4420 |
| ASV19 | *Bacteroides* | 15 | 0 |
| ASV22 | *Alcaligenes* | 708 | 693 |
| ASV24 | *Aquamicrobium* | 545 | 501 |
| ASV25 | *Bacteroides* | 0 | 11 |
| ASV33 | *Peptostreptococcus* | 0 | 7 |
| ASV47 | *UCG-002* | 6 | 0 |
| ASV65 | *Clostridium sensu stricto 1* | 8 | 0 |
| ASV83 | *Bilophila* | 0 | 7 |
| ASV106 | *Cutibacterium* | 0 | 87 |
| ASV119 | *Lawsonella* | 14 | 98 |
| ASV132 | *Sutterella* | 5 | 0 |
| ASV140 | *Staphylococcus* | 66 | 57 |
| ASV174 | *Clostridium sensu stricto 1* | 21 | 0 |
| ASV179 | *Arthrobacter* | 32 | 13 |
| ASV203 | *Enhydrobacter* | 6 | 14 |
| ASV225 | *Microbacterium* | 0 | 92 |
| ASV237 | *Rhodococcus* | 39 | 24 |
| ASV275 | *Corynebacterium* | 0 | 41 |
| ASV277 | *Staphylococcus* | 0 | 55 |
| ASV289 | *Cutibacterium* | 0 | 19 |
| ASV310 | *Microlunatus* | 33 | 0 |

| | | | |
|---|---|---|---|
| ASV319 | *Brevibacterium* | 0 | 29 |
| ASV341 | *Chryseobacterium* | 44 | 0 |
| ASV344 | *Rothia* | 0 | 8 |
| ASV369 | *NS5 marine group* | 11 | 0 |
| ASV375 | *Anaerococcus* | 0 | 9 |
| ASV410 | *Pelomonas* | 7 | 6 |
| ASV427 | *Lentibacter* | 33 | 0 |
| ASV445 | *Microbacterium* | 31 | 0 |
| ASV503 | *Clade Ia* | 26 | 0 |
| ASV522 | *Weeksellaceae Unclassified* | 25 | 0 |
| ASV523 | *AUTHM297* | 25 | 0 |
| ASV524 | *Dolosigranulum* | 20 | 0 |
| ASV555 | *Caulobacter* | 0 | 3 |
| ASV559 | *Flavobacterium* | 23 | 0 |
| ASV597 | *Bradyrhizobium* | 7 | 0 |
| ASV615 | *Staphylococcus* | 20 | 0 |
| ASV640 | *Reyranella* | 15 | 0 |
| ASV644 | *Altererythrobacter* | 19 | 0 |
| ASV662 | *Paracoccus* | 0 | 18 |
| ASV663 | *Microbacterium* | 0 | 18 |
| ASV687 | *Brevundimonas* | 17 | 0 |
| ASV722 | *Deinococcus* | 16 | 0 |
| ASV754 | *Brachybacterium* | 0 | 6 |
| ASV767 | *Microbacteriaceae Unclassified* | 0 | 15 |
| ASV796 | *Streptococcus* | 0 | 8 |
| ASV865 | *Staphylococcus* | 13 | 0 |
| ASV932 | *Granulicatella* | 12 | 0 |

| | | | |
|---|---|---|---|
| ASV933 | *Parasegetibacter* | 0 | 12 |
| ASV934 | *Gemmobacter* | 0 | 12 |
| ASV1015 | *Chroococcidiopsaceae Unclassified* | 11 | 0 |
| ASV1106 | *Flavobacteriaceae Unclassified* | 10 | 0 |
| ASV1195 | *Corynebacterium* | 9 | 0 |
| ASV1196 | *Campylobacter* | 0 | 9 |
| ASV1316 | *Alloprevotella* | 8 | 0 |
| ASV1317 | *Novosphingobium* | 0 | 8 |
| ASV1318 | *Blastococcus* | 0 | 6 |
| ASV1464 | *67-14 Unclassified* | 7 | 0 |
| ASV1465 | *0319-6G20 Unclassified* | 7 | 0 |
| ASV1466 | *CHKCI001* | 7 | 0 |
| ASV1467 | *Lactobacillus* | 0 | 7 |
| ASV1468 | *Carnobacterium* | 0 | 7 |
| ASV1469 | *Flavobacterium* | 0 | 7 |
| ASV1470 | *CK06-06-Mud-MAS4B-21* | 0 | 7 |
| ASV1647 | *Craurococcus-Caldovatus* | 0 | 6 |
| ASV1648 | *Leptotrichia* | 0 | 6 |
| ASV1649 | *Streptococcus* | 0 | 6 |
| ASV1815 | *Lachnospiraceae Unclassified* | 5 | 0 |
| ASV1816 | *Paenibacillus* | 5 | 0 |
| ASV1817 | *Corynebacterium* | 5 | 0 |
| ASV1818 | *Pseudorhodobacter* | 0 | 5 |
| ASV1819 | *Saccharimonadales Unclassified* | 0 | 5 |
| ASV1991 | *Acinetobacter* | 4 | 0 |
| ASV1992 | *Gaiellales Unclassified* | 0 | 4 |
| ASV2121 | *Paracoccus* | 3 | 0 |

| | | | |
|---|---|---|---|
| ASV2122 | *Lachnospiraceae Unclassified* | 3 | 0 |
| ASV2123 | *Nocardioides* | 0 | 3 |
| ASV2124 | *Actinomyces* | 0 | 3 |
| ASV2208 | *Spirosoma* | 2 | 0 |
| ASV2209 | *DSSD61* | 0 | 2 |

**Table S7**: Contaminant reads and ASVs were detected in aPBS blanks (n=2 biological replicates) using Illumina V3-V4 amplicon 16S rRNA gene sequencing.

| **ASV** | **Genus** | **Read counts of replicates** | |
|---|---|---|---|
| ASV10 | *Streptococcus* | 9 | 0 |
| ASV13 | *Cutibacterium* | 593 | 369 |
| ASV15 | *Fusobacterium* | 9 | 0 |
| ASV52 | *Veillonella* | 2 | 0 |
| ASV61 | *Clostridium sensu stricto 1* | 16 | 0 |
| ASV62 | *Alloprevotella* | 12 | 0 |
| ASV88 | *Rothia* | 8 | 0 |
| ASV112 | *Bacteroides* | 13 | 0 |
| ASV129 | *Alloprevotella* | 10 | 0 |
| ASV149 | *UCG-002* | 7 | 0 |
| ASV153 | *Lachnoanaerobaculum* | 8 | 0 |
| ASV171 | *[Ruminococcus] torques group* | 13 | 0 |
| ASV172 | *Faecalibacterium* | 10 | 0 |
| ASV179 | *Arthrobacter* | 23 | 19 |
| ASV203 | *Enhydrobacter* | 6 | 0 |
| ASV206 | *UCG-005* | 5 | 0 |
| ASV263 | *Alysiella* | 5 | 0 |
| ASV275 | *Corynebacterium* | 0 | 15 |
| ASV277 | *Staphylococcus* | 0 | 10 |
| ASV310 | *Microlunatus* | 12 | 0 |
| ASV378 | *Cyanobium PCC-6307* | 13 | 8 |
| ASV400 | *CL500-29 marine group* | 0 | 6 |
| ASV411 | *Prevotella_7* | 35 | 0 |
| ASV555 | *Caulobacter* | 12 | 0 |

| | | | |
|---|---|---|---|
| ASV556 | *Brevundimonas* | 0 | 11 |
| ASV572 | *Friedmanniella* | 22 | 0 |
| ASV593 | *Sphingomonas* | 14 | 0 |
| ASV600 | *Brevibacterium* | 0 | 8 |
| ASV606 | *NS3a marine group* | 0 | 11 |
| ASV641 | *Candidatus Aquiluna* | 13 | 0 |
| ASV658 | *Rhodoplanes* | 6 | 0 |
| ASV858 | *Paracoccus* | 4 | 0 |
| ASV868 | *RS62 marine group* | 13 | 0 |
| ASV1018 | *Chthoniobacter* | 11 | 0 |
| ASV1114 | *Ruminococcus* | 10 | 0 |
| ASV1115 | *Brachybacterium* | 0 | 10 |
| ASV1202 | *Prevotellaceae Unclassified* | 9 | 0 |
| ASV1203 | *CL500-29 marine group* | 0 | 9 |
| ASV1325 | *Cellvibrio* | 8 | 0 |
| ASV1326 | *Nocardioides* | 8 | 0 |
| ASV1327 | *Sphingomonas* | 0 | 8 |
| ASV1328 | *Knoellia* | 0 | 8 |
| ASV1476 | *Rhodobacteraceae Unclassified* | 7 | 0 |
| ASV1655 | *Sphingopyxis* | 6 | 0 |
| ASV1656 | *Deinococcus* | 6 | 0 |
| ASV1657 | *Vicinamibacterales Unclassified* | 6 | 0 |
| ASV1658 | *Corynebacterium* | 0 | 6 |
| ASV1825 | *EF100-94H03 Unclassified* | 5 | 0 |
| ASV1826 | *Aerococcus* | 5 | 0 |
| ASV1827 | *Brachybacterium* | 5 | 0 |
| ASV1828 | *Jonquetella* | 0 | 5 |

| | | | |
|---|---|---|---|
| ASV1829 | *Microbacterium* | 0 | 5 |
| ASV1830 | *Paracoccus* | 0 | 5 |
| ASV1999 | *Corynebacterium* | 4 | 0 |
| ASV2000 | *Lachnospiraceae Unclassified* | 4 | 0 |
| ASV2001 | *Frankiales Unclassified* | 0 | 4 |

**Table S8**: Contaminant reads and ASVs were detected in the sorted mock cells (n=3 biological replicates) and noise (n=2 biological replicates) using Illumina V3-V4 amplicon 16S rRNA gene sequencing.

| ASV | Genus | Sorted cells (n=3) | | | Sorted noise (n=2) | |
|---|---|---|---|---|---|---|
| | | Reads | Reads | Reads | Reads | Reads |
| ASV11 | *Pseudomonas* | 530 | 400 | 0 | 1658 | 71 |
| ASV13 | *Cutibacterium* | 89 | 107 | 0 | 86 | 0 |
| ASV15 | *Fusobacterium* | 0 | 11 | 0 | 0 | 0 |
| ASV22 | *Alcaligenes* | 116 | 177 | 0 | 366 | 13 |
| ASV24 | *Aquamicrobium* | 152 | 123 | 0 | 262 | 10 |
| ASV237 | *Rhodococcus* | 0 | 0 | 0 | 7 | 0 |
| ASV375 | *Anaerococcus* | 0 | 0 | 0 | 11 | 0 |
| ASV518 | *PeM15 Unclassified* | 0 | 0 | 0 | 25 | 0 |
| ASV556 | *Brevundimonas* | 0 | 7 | 0 | 0 | 0 |
| ASV593 | *Sphingomonas* | 0 | 0 | 0 | 7 | 0 |
| ASV640 | *Reyranella* | 4 | 0 | 0 | 0 | 0 |
| ASV641 | *Candidatus Aquiluna* | 0 | 6 | 0 | 0 | 0 |
| ASV810 | *hgcI clade* | 0 | 14 | 0 | 0 | 0 |
| ASV925 | *Prevotella_7* | 12 | 0 | 0 | 0 | 0 |
| ASV1074 | *Phenylobacterium* | 5 | 0 | 0 | 0 | 0 |
| ASV1434 | *Bacillaceae Unclassified* | 0 | 4 | 0 | 0 | 0 |
| ASV1629 | *Massilia* | 0 | 6 | 0 | 0 | 0 |
| ASV1800 | *Mitochondria Unclassified* | 5 | 0 | 0 | 0 | 0 |
| ASV1801 | *Rubrobacter* | 0 | 0 | 0 | 5 | 0 |
| ASV1802 | *Microbacteriaceae Unclassified* | 0 | 0 | 0 | 5 | 0 |
| ASV2114 | *Crocinitomix* | 0 | 3 | 0 | 0 | 0 |

**Table S9**: *Veillonella atypica* (Rogosa 1965) relic DNA, spiked into a ZymoBIOMICS microbial community standard (mock) and PBS, was depleted in sorted cells ($10^5$ cells $mL^{-1}$) and emerged in sorted noise ($10^5$ noise events $mL^{-1}$). Benzonase nuclease activity selectively depleted relic DNA of *Veillonella atypica, Pseudomonas aeruginosa* ASV63, *Escherichia-Shigella* coli ASV 11/42/43/77/88/179 and *Salmonella enterica* ASV 25/201. AMPure XP clean-up eliminated relic DNA but was not selective resulting in a distorted community composition. A decrease in the (relic) DNA concentration prior to sorting resulted in an inflation in absolute and relative contaminant sequencing read counts attesting the need of *in silico* data decontamination. Samples were sequenced with PacBio full-length 16S rRNA gene sequencing. Cells were sorted using a non-selective SYBR® Green I staining. A FACS sheath fluid blank was included to identify and track the FACSome contaminants introduced during FACS-sequencing.

| **Community** | **Fraction** | **Treatment** | ***V.atypica* reads** | **% *V.atypica*** | **Contaminant reads** | **Total reads** | **DNA concentration (pg/µL)** |
|---|---|---|---|---|---|---|---|
| theory | | none | | 9.3 | | | |
| mock | cell | untreated | 89 | 0.3 | 264 | 28818 | 19.2 |
| mock | noise | untreated | 238 | 5.8 | 589 | 4097 | 0.6 |
| PBS | noise | untreated | 1018 | 47.8 | 1312 | 2132 | 5.1 |
| PBS | noise | untreated | 252 | 17.0 | 489 | 1482 | 3.6 |
| mock | cell | Benzonase | 240 | 2.4 | 965 | 9868 | 22.2 |
| mock | noise | Benzonase | 31 | 1.4 | 167 | 2147 | 0.4 |
| PBS | noise | Benzonase | 19 | 0.3 | 3335 | 6108 | 5.2 |
| PBS | noise | Benzonase | 0 | 0.0 | 3689 | 6639 | 34.6 |
| mock | cell | AMPure | 40 | 0.4 | 2556 | 9163 | 30.4 |
| mock | noise | AMPure | 27 | 0.5 | 2843 | 5963 | 0.9 |
| PBS | noise | AMPure | 0 | 0.0 | 2492 | 3672 | 2.5 |
| PBS | noise | AMPure | 131 | 3.0 | 2465 | 4312 | 23.4 |
| sheath | noise | none | 0 | 0.0 | 2120 | 3231 | 62.8 |
